# Crowdsourced analysis of ash and ash dieback through the Open Ash Dieback project: A year 1 report on datasets and analyses contributed by a self-organising community

**DOI:** 10.1101/004564

**Authors:** Diane Saunders, Kentaro Yoshida, Christine Sambles, Rachel Glover, Bernardo Clavijo, Manuel Corpas, Daniel Bunting, Suomeng Dong, Ghanasyam Rallapalli, Matthew D. Clark, David Swarbreck, Sarah Ayling, Matthew Bashton, Steve Collin, Tsuyoshi Hosoya, Anne Edwards, Lisa Crossman, Graham Etherington, Joe Win, Liliana Cano, David J. Studholme, J. Allan Downie, Mario Caccamo, Sophien Kamoun, Dan MacLean

## Abstract

Ash dieback is a fungal disease of ash trees caused by *Hymenoscyphus pseudoalbidus* that has swept across Europe in the last two decades and is a significant threat to the ash population. This emergent pathogen has been relatively poorly studied and little is known about its genetic make-up. In response to the arrival of this dangerous pathogen in the UK we took the unusual step of providing an open access database and initial sequence datasets to the scientific community for analysis prior to performing an analysis of our own. Our goal was to crowdsource genomic and other analyses and create a community analysing this pathogen. In this report on the evolution of the community and data and analysis obtained in the first year of this activity, we describe the nature and the volume of the contributions and reveal some preliminary insights into the genome and biology of *H. pseudoalbidus* that emerged. In particular our nascent community generated a first-pass genome assembly containing abundant collapsed AT-rich repeats indicating a typically complex genome structure. Our open science and crowdsourcing effort has brought a wealth of new knowledge about this emergent pathogen within a short time-frame. Our community endeavour highlights the positive impact that open, collaborative approaches can have on fast, responsive modern science.

## 1 Introduction

Chalara dieback of ash is caused by the aggressive fungal pathogen *Hymenoscyphus pseudoalbidus* (anamorph *Chalara fraxinea*), an emergent disease of ash trees (*Fraxinus excelsior*). It causes a range of symptoms that include dieback of tree crowns, wilting leaves, lesions and cankers on leaves, stems and roots, dieback of leaves, browning of petioles and staining and damage to woody tissues beneath lesions [1]. The disease can be fatal for young saplings, whilst providing opportunities for secondary infection in mature trees ultimately killing or severely inhibiting their growth. In Denmark the infection of up to 90% of the ash tree population has been attributed to ash dieback.

With this in mind, up to 90% of the more than 80 million ash trees in the UK are thought to be under threat. The disease, which is a newcomer to Britain, was first reported in the natural environment in October 2012 and has since been recorded in native woodland throughout the UK. There is no known treatment for Chalara dieback of ash, current control measures include burning infected trees to try and prevent spread. Upon its discovery there was a large public and media outcry and legal challenges forcing the British government into taking fast action [2].

To kick start genomic analyses of the pathogen and host, we took the unconventional step of rapidly generating and releasing genomic sequence data prior to any analysis. We released the data through our ash and ash dieback website, http://oadb.tsl.ac.uk [3], which we launched in December 2012. Speed is essential in response to emerging and severely threatening diseases. By publicly releasing genomic data we aimed to make it possible for experts from around the world to come together and assist in the analysis, rapidly accelerating the process.

Our activity stimulated a community to come together and act rapidly and openly. Here we will bring together and discuss the progress of this unusual project and highlight some of the key findings about *H. pseudoalbidus* that emerged from this exercise as a direct result of crowdsourcing genomic analysis from scientists.

## 2 Creation of a community hub for crowdsourced analysis

### 2.1 OpenAshDieBack - a news and report hub

Our primary hub is OpenAshDieBack (OADB), a website at http://oadb.tsl.ac.uk, through which announcements of new datasets can be made and also through which analyses can be presented. As it is primarily a news and article website, essentially a large blog, OADB provides a friendly and intuitive way to read news from the project as it arrives and is a convenient way to illustrate the outputs from the community. Contribution to this site is done by submission of text to the maintainers via email to, who will then post the submission.

Data is not held directly on OADB since a news website is not easily compatible with holding structured data for general access. As such we link to a git based database hosted on GitHub at https://github.com/ash-dieback-crowdsource/data.

### 2.2 ash-dieback-crowdsource - an open access GitHub repository database

For crowdsourcing we need to be able to track atomic contributions from lots of contributors, automatically, reliably and as they happen. The version control software git, and its associated hosting platform GitHub is a system that provides such functionality. Git and GitHub were conceived as tools for computer programmers who typically work in widely distributed groups on exactly the same set of files. In practice a contributor starts the day by pulling the latest version of the data from GitHub to a local machine, makes the changes/additions/new data that they think are appropriate and pushes the files back for integration. Git sits in the middle keeping track of what changes were made and who made them and provides a framework for merging any conflicts.

In order to share data and make contributions clearly attributable to the provider we set up a public repository on the GitHub website at https://github.com/ash-dieback-crowdsource/data. This was inspired by the earlier *Escherichia coli* 0104:H4 repository [4], but with a richer, contextual structure and stronger guidelines and provision for meta-data definition.

Our repository has a generic data structure designed to hold multiple sets of genomics data and links to external sources of data too large to store on GitHub, such as raw sequence reads. The repository is essentially a common directory tree (folder-based) structure, semantically organised, so that a user should be able to follow and find what is needed intuitively. The base folder contains the project level information and subdivides through organism and strain folders to data folders, named by data type and containing actual datasets and meta-data descriptions. With this data structure, each dataset has a unique, rule-based and logical address (see Figure 1 for an example).

**Figure 1:**
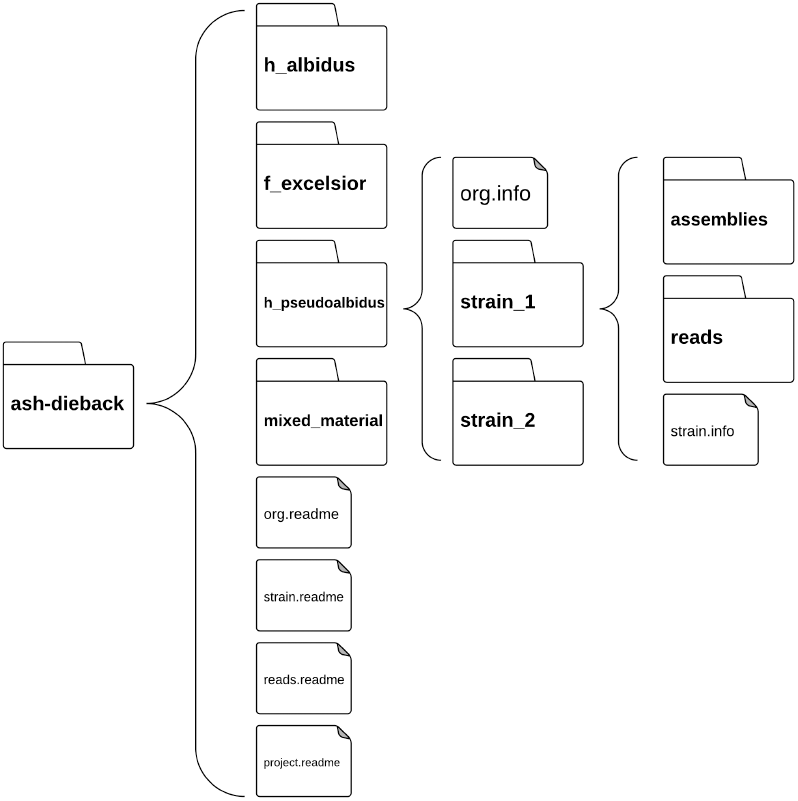
Example of the basic folder structure of the Ash Dieback GitHub repository. Folder icons represent folders on the file system, open braces indicate contents of folders. The top level (leftmost) folder contains the project and can be followed down through the organisms to the data specific to each organism and ultimately strain. At each level files suffixed .info or .readme contain or define the metadata of the data in the repository. Original available at http://dx.doi.org/10.6084/m9.figshare.1004999

Provision for meta-data is made with files suffixed .README and .info. The .README file kept in the root folder define the meta-data required for data-types lower in the structure while actual meta-data is recorded in .info files. Each .info describes one discrete dataset contribution.

### 2.3 Activity in the repository

The GitHub ash-dieback organisation on GitHub currently has 43 members from institutes around the world, including the United Kingdom, France, Canada and the United States. As of the 30th November 2013, the repository has received 170 contributions (measured as commits to the repository) from 13 of those 43 members. Multiple members of the same physical lab group may have coordinated submission via a single member, potentially underestimating the actual numbers of contributors.

Although not all commits are novel contributions, commit activity can be a useful proxy metric for measuring contributions to the repository. The pattern of total commits by time and by volume is represented in Figure 2. The initial months saw the highest volume, with over 35 million lines of data added of which 10 million were removed, or edited. This flurry of activity represents deposition of the first few datasets and the ripe opportunity in which the most tractable analyses were carried out. Throughout the rest of the year there was a less massive but still significant contribution continuing at intervals. The periodic pattern is attributable to the greater computation and human-effort required for bigger and more complex analysis that were done later. A full time-line of commits can be seen in Supplemental Table 1 and major data sets and analysis contributions in the database are outlined in Figure 3.

**Figure 2:**
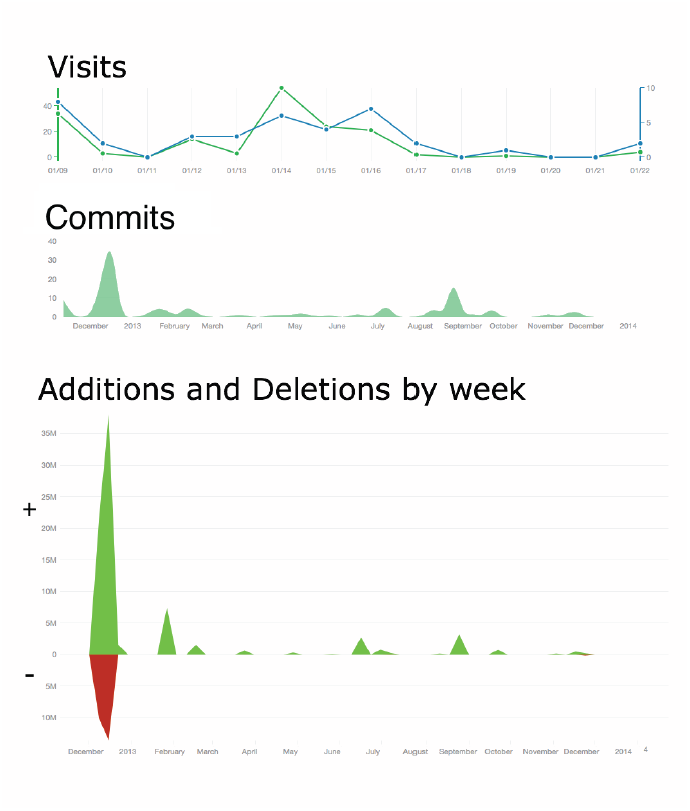
Activity on the database. Top: GitHub Analytics representing unique and total page views for URL https://github.com/ash-dieback-crowdsource/data/ in the first possible reporting period for this website. GitHub released analytics on 7th January, 2014, so data are presented from that date to 22nd January 2014. Middle: Commit events over the whole period of the project to from December 16, 2012 to January 22, 2014. Bottom: Individual lines of data added (green areas) or deleted (red areas) over the period December 16 2012 to January 22, 2014. Original available at http://dx.doi.org/10.6084/m9.figshare.1005000

**Figure 3:**
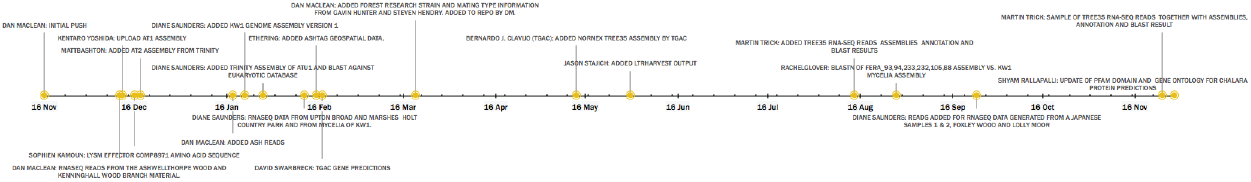
Major commit events to the database over the first twelve months. Original version available at http://dx.doi.org/10.6084/m9.figshare.1005001

Use of the data without contribution is not directly tracked, reports like this are not available from GitHub. Web access figures (analogous to Google Analytics) to the GitHub repository have been available only since GitHub began reporting figures to it’s user base on 7th January 2014. From this point to the 22nd January the repository received a total of 174 views from 37 unique addresses (see Figure 2 for profile), indicating access and interest in the data in a traditionally quiet period. It is unwise to extrapolate from such a small sample over a longer period but it is encouraging that so many accesses were made in the new year and suggests steady interest in the data and a useful resource. The growth of the repository and website indicates that OADB is providing a useful and well-accessed database and news site for the fledgling ash dieback community.

OpenAshDieBack contains a wide range of datasets contributed through crowdsourcing including genomic DNA sequence and assemblies for *H.pseudoalbidus* RNA-Seq data and transcriptome assemblies from multiple strains, numerous annotations on these large data sets and multiple images of the morphology and reports on the pathology of the fungus.

The 30 genomic and RNA-Seq datasets mentioned in this report are summarised in Table 1, which describes the organism or material type from which they were collected, the data type and a short name and the location from which they were collected, if recorded. The majority are RNA-seq datasets from isolated cultures of *H.pseudoalbidus* or samplings of mixed material (a sample of infected wood that yields a metagenomic sample when DNA or RNA is extracted). The analyses of these data sets are presented below.

**Table 1:** Sequence read and assembly datasets available in OADB. Full text version available at http://dx.doi.org/10.6084/m9.figshare.1005036

| Organism | Tissue | Data Type | Short Name | Location Collected |
| --- | --- | --- | --- | --- |
| H.pseudoalbidus | Mature Fruiting Body | RNA Seq Assembly | MFB1 |  |
| H.pseudoalbidus | Mature Fruiting Body | RNA Seq Reads | MFB1 |  |
| H.pseudoalbidus | Primordial Fruiting Body | RNA Seq Assembly | PFB1 |  |
| H.pseudoalbidus | Primordial Fruiting Body | RNA Seq Reads | PFB1 |  |
| Fraxinus excelsior | Uninfected Branch Pith | RNA Seq Reads | ATU1 | Ashwellthorpe, Norfolk |
| Fraxinus excelsior | Uninfected Branch Pith | RNA Seq Assembly | ATU1 | Ashwellthorpe, Norfolk |
| H.pseudoalbidus | Isolated Culture | gDNA Reads | KW1 | Kenninghall Wood, Norfolk |
| H.pseudoalbidus | Isolated Culture | RNA Seq Reads |  |  |
| H.pseudoalbidus | Isolated Culture | gDNA Assembly | KW1 | Kenninghall Wood, Norfolk |
| H.pseudoalbidus | Isolated Culture | RNA Seq Assembly |  |  |
| H.pseudoalbidus | Isolated Culture | RNA Seq Reads | FERA_105 | Green Ln, Lyminge, Kent |
| H.pseudoalbidus | Isolated Culture | RNA Seq Reads | FERA_232 | Cronkhill Ln, Carlton, Yorks |
| H.pseudoalbidus | Isolated Culture | RNA Seq Reads | FERA_233 | Cronkhill Ln, Carlton, Yorks |
| H.pseudoalbidus | Isolated Culture | RNA Seq Reads | FERA_88 | Seaton Rd, Hornsea, Yorks |
| H.pseudoalbidus | Isolated Culture | RNA Seq Reads | FERA_93 | Thetford Rd, Nr Wretham, Norfolk |
| H.pseudoalbidus | Isolated Culture | RNA Seq Reads | FERA_94 | Thetford Rd, Nr Wretham, Norfolk |
| H.pseudoalbidus | Isolated Culture | RNA Seq Assembly | FERA_105 |  |
| H.pseudoalbidus | Isolated Culture | RNA Seq Assembly | FERA_232 |  |
| H.pseudoalbidus | Isolated Culture | RNA Seq Assembly | FERA_233 |  |
| H.pseudoalbidus | Isolated Culture | RNA Seq Assembly | FERA_88 |  |
| H.pseudoalbidus | Isolated Culture | RNA Seq Assembly | FERA_93 |  |
| H.pseudoalbidus | Isolated Culture | RNA Seq Assembly | FERA_94 |  |
| H.pseudoalbidus | Mixed Material | RNA Seq Reads | FW1 | Foxley Wood, Norfolk |
| H.pseudoalbidus | Mixed Material | RNA Seq Reads | LM1 | Lolly Moor, Norfolk |
| H.pseudoalbidus | Mixed Material | RNA Seq Reads | JP1 |  |
| H.pseudoalbidus | Mixed Material | RNA Seq Reads | JP2 |  |
| H.pseudoalbidus | Mixed Material | RNA Seq Reads | UB1 | Upton Broad and Marshes |
| H.pseudoalbidus | Mixed Material | RNA Seq Reads | AT1 | Ashwellthorpe, Norfolk |
| H.pseudoalbidus | Mixed Material | RNA Seq Reads | AT2 | Ashwellthorpe, Norfolk |
| H.pseudoalbidus | Mixed Material | RNA Seq Reads | HC1 | Holt Country Park, Norfolk |

## 3 Genome Architecture and Content of *Hymenoscyphus pseudoalbidus*

### 3.1 The KW1 draft assembly of the *H. pseudoalbidus* genome

An isolate of *Hymenoscyphus pseudoalbidus* was collected and isolated from a tree in Kenninghall Wood, Norfolk, UK, a site close to Ashwellthorpe Lower Wood where ash dieback was first noted in the wild in the United Kingdom. DNA was extracted from a purified culture and sequenced according to the procedure described in section 10.1. Genomic DNA was sequenced using mate-pair libraries with insert sizes of ~196 nt and ~570 nt. An assembly was constructed as described in methods and immediately deposited in the OADB GitHub. Assembly statistics such as NG(X) were calculated as described in section 10.2. The estimated genome size of *H.pseudoalbidus* KW1 is approximately 63 Mbp and the total assembly length covers 63,153,926 bp in just 1672 scaffolds. We have no prior or independent information on the size of the *H.pseudoalbidus* genome and this is our only estimate. The distribution of scaffold length shows consistently large contigs, the NG(50) [5] is long at 70,504 bp and the distribution of sequence lengths shows that over 96% of the entire assembly is in scaffolds of greater than 10 kbp in length (see Figure 4). As our size estimate is based on this assembly then we cannot claim that the assembly covers any particular proportion of the genome as is usually done with an NG(50). However, NG(50) remains useful for qualitatively describing our assembly.

**Figure 4:**
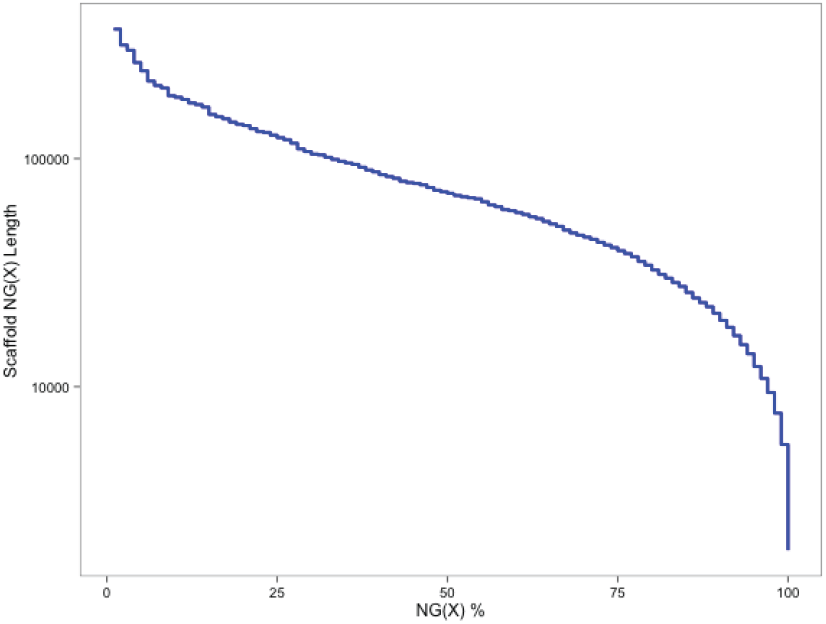
NG graph showing an overview of scaffold lengths in the Kenninghall Wood (KW1) genomic DNA assembly. Scaffold lengths are sorted in greatest to shortest order and summed until that sum exceeds the assembly length percentile for all integer percent values between 1 % and 100 %. The Scaffold NG(X) Length is taken as the length of the scaffold that makes the descending length sum of scaffolds exceed the Xth percentile of the current expected assembly size of 63 Mbp. We do not have another independent size estimate for this genome. Original available at http://dx.doi.org/10.6084/m9.figshare.1005002

We assessed the gene content of the assembly using CEGMA [6] to evaluate the presence of 248 core conserved eukaryotic genes (see methods 2). This analysis showed 236 of the 248 (95%) of the ultra-conserved core genes are present in the assembly (Table 2) indicating excellent reconstruction of the gene space. Taken together with the statistics above we conclude that the KW1 draft assembly is useful for downstream analyses.

**Table 2:** Results from CEGMA [6] analysis of the KW1 scaffold assembly assessing completeness of conserved core eukaryotic genes in the sequence.

|  | Number of Proteins | Percent Completeness | Total | Average | Over 1 Orthologue |
| --- | --- | --- | --- | --- | --- |
| Complete | 236 | 95.16 | 260 | 1.10 | 8.05 |
| Group 1 | 61 | 92.42 | 66 | 1.08 | 6.56 |
| Group 2 | 53 | 94.64 | 60 | 1.13 | 9.43 |
| Group 3 | 59 | 96.72 | 69 | 1.17 | 13.56 |
| Group 4 | 63 | 96.92 | 65 | 1.03 | 3.17 |
| Partial | 241 | 97.18 | 274 | 1.14 | 10.37 |
| Group 1 | 63 | 95.45 | 69 | 1.10 | 7.94 |
| Group 2 | 54 | 96.43 | 63 | 1.17 | 12.96 |
| Group 3 | 60 | 98.36 | 74 | 1.23 | 16.67 |
| Group 4 | 64 | 98.46 | 68 | 1.06 | 4.69 |

Annotation *ab initio*, with cross-referenced protein evidence from four other sequenced fungi and using RNA-seq datasets was carried out as described in section 10.3. The combined KW1 annotation contains 10,252 separate gene models (see Table 6), a total similar to that in other sequenced Leotiomycetes [7].

### 3.2 Variant analysis of different isolates and fruiting bodies of *H.pseudoalbidus*

To identify the extent of genetic variation in the sequenced samples, we undertook variant analysis from seven isolates (KW1, FERA 105, FERA 232, FERA 233, FERA 88, FERA 93 and FERA 94), two fruiting body samples (MFB1 and PFB1) and four mixed material samples (HC1, UB1, AT1 and AT2) as described in section 10.17. The FERA samples contain fewer sequencing reads compared to the fruiting body samples, mixed material samples and KW1 (Table 4). Sample KW1 covers 32% of the genome positions with a coverage of >10X compared to between 0.17-0.27% for the FERA samples. The remaining samples range from 13% (AT1) to 32% (UB1) (see Table 4). This suggests that limited SNP calling is possible in the FERA samples.

SNP analysis as described in section 10.17 revealed a high level of genetic variation between the samples (Table 5). The mature fruiting body (MFB1) and primordial fruiting body (PFB1) samples showed a high level of variation (approx. 16,000 SNPs) when aligned to the KW1 genome. Samples AT1 (approx. 6,000), AT2 (approx. 3,000), HB1 (approx. 14,000) and UB1 (approx. 12,000) also suggested a high level of genetic variation although this could be due to the alignment of non-*H.pseudoalbidus* reads present in the mixed sample, which have a high degree of similarity to regions of the *H.pseudoalbidus* genome. This highlights the importance of the availability of multiple genomes.

Further analysis is required to understand the significance of this difference in variation and whether the UB1 and HC1 variation is due to the presence of other fungi, such as *Phytothphora* sp. and *Togninia* sp. identified in the mixed samples interfering with the SNP calling.

### 3.3 The genome of *Hymenoscyphus pseudoalbidus* KW1 harbours many AT-rich repeat islands that are not yet assembled fully

The genomes of many filamentous plant pathogens have been shown to be rich in repeats and transposable elements [8]. In particular many of these regions appear to be enriched in elements and genes that can be shown to be under greater selective pressure than elements in other regions of the genome, such as genes encoding disease inducing effector proteins. Effector proteins are key to the armoury of the pathogen, acting within the host plant to manipulate host physiology in favour of pathogen progression. Therefore it is essential to assess genome architecture and its contribution to directional selection in particular genomic regions. To examine any collapsed structure of the genome from the draft level assembly we re-aligned the genomic DNA reads used to create the assembly of the KW1 isolate to assess coverage and mate-pair distance distributions (section 10.9). We compared this with GC-content and gene and repeat feature density and discovered a striking pattern of AT-rich regions in which nucleotide content of the reference sequence dropped below 30% GC in long stretches (see Figures 5 and 6). The positions of these are uniformly co-localised with the regions in which reads aligned back to the assembly are vastly more numerous relative to the rest of the genome. Furthermore, peaks of abundance of re-aligned reads co-localise with troughs of GC content. Repeat sequence searches using the RepeatMasker database and transposable element analysis also show peaks of abundance of matches coincident with the AT-rich, high-coverage areas. Finer grained analysis of the GC content of individual feature types from the annotation of the assembly described in section 10.3 revealed that coding features and Illumina RNA-seq reads are not AT-rich and that this is a property of repeat and intron sequences (Figure 7).

**Figure 5:**
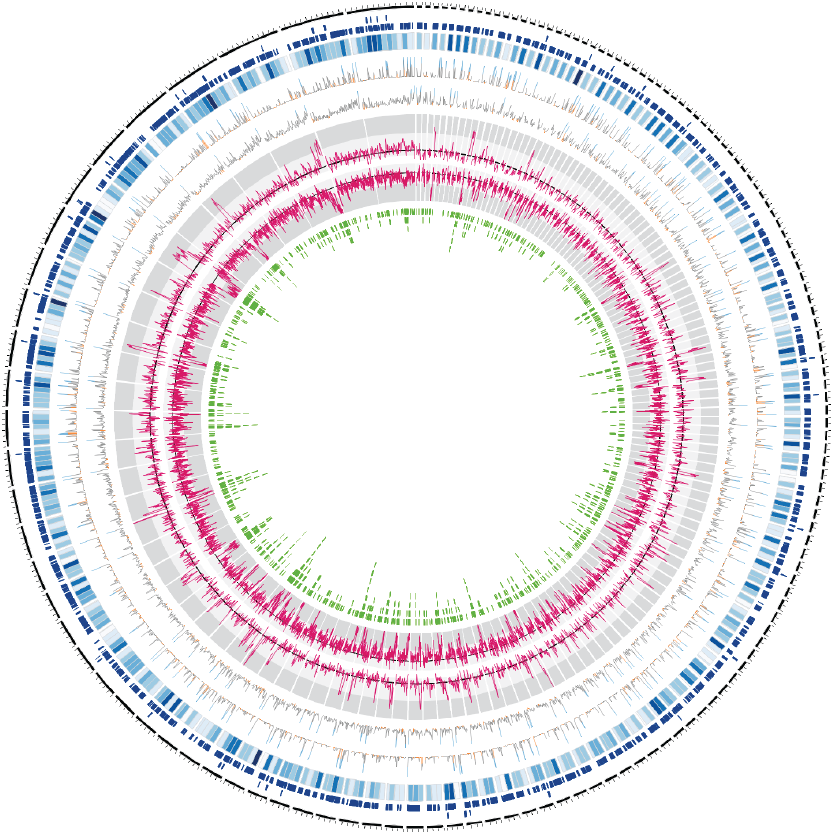
The KW1 assembly of *Hymenoscyphus pseudoalbidus* displaying: outer black ring - scaffolds of length of at least 10 kbp; blue stacks - gene models; blue heatmap - gene density in 10 kbp windows; line plot - aligned distance between paired end read for library with 196 bp insert size (<31 bp - 5th percentile - rendered in orange, >351 bp - 95th percentile - rendered in blue);line plot aligned distance between paired end read for library with 570 bp insert size (<265 bp - 5th percentile - rendered in orange, *>*2313 bp - 95th percentile - rendered in blue) Thresholds calculated as per https://github.com/danmaclean/h_pseu_analysis/blob/master/circos/scripts/assess_insert_size_distributions.md pink line plot - uniquely mapped read coverage. Maximum plotted = 300, minimum plotted = 80; pink line plot - GC percent, black line = 50% GC, light grey area 50% - 30% GC, dark grey area *>*30% GC. Maximum plotted = 60% GC, minimum plotted = 20% GC ; green stacks - Repeat Masker matches. Original available at http://dx.doi.org/10.6084/m9.figshare.791640

**Figure 6:**
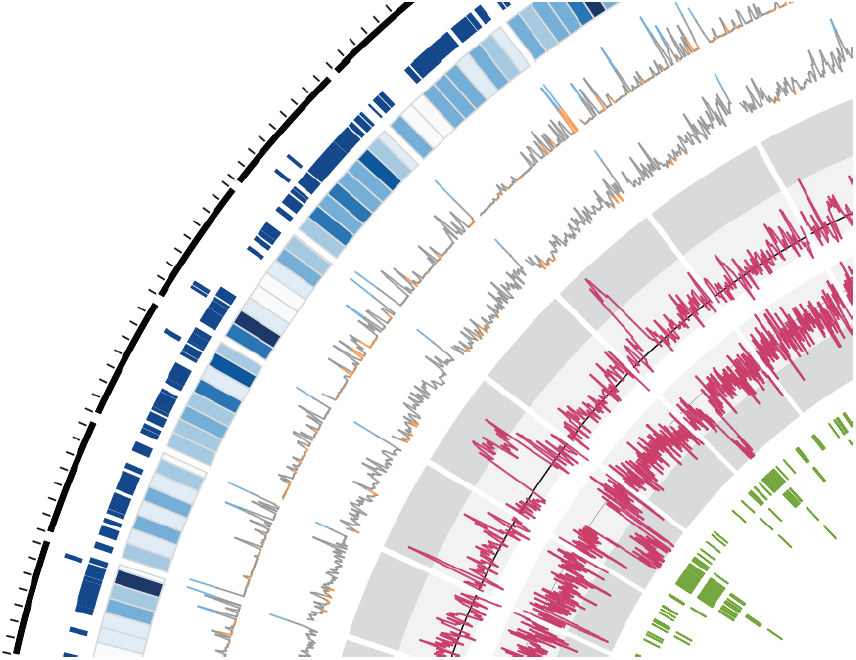
Zoom region of the KW1 assembly of *Hymenoscyphus pseudoalbidus* displaying tracks as per Figure 5. Original available at http://dx.doi.org/10.6084/m9.figshare.791640

**Figure 7:**
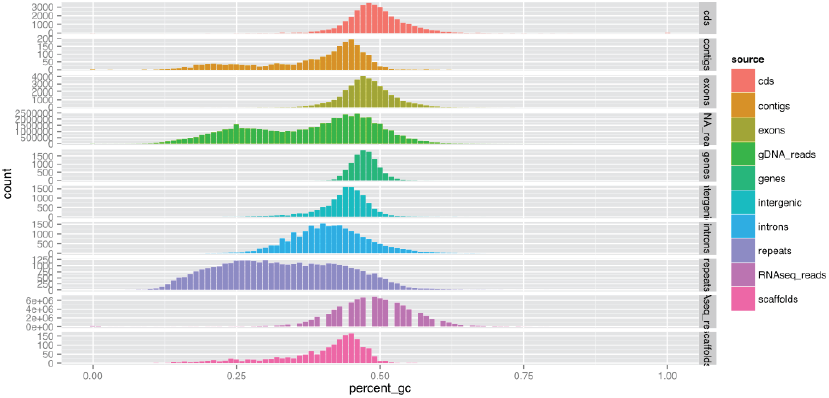
Distribution of KW1 genome assembly GC content in different genomic annotation feature and assembly types and raw sequence reads. Original available at http://dx.doi.org/10.6084/m9.figshare.706771

Together these analyses suggest numerous small collapsed regions in the assembly, a minor issue of the snapshot state of the assembly which is unavoidable and in fact somewhat expected with the sequence libraries used. An artefact like this can easily be corrected with the inclusion of longer reads and insert sizes as is planned in the next phase of sequencing. Notwithstanding, the current version of the assembly has provided us with an exciting hypothesis about the structure of the *H.pseudoalbidus* genome: it has many repeat-rich, potentially adaptable genome regions that could contribute to extreme genomic plasticity and as such may provide the raw material for evolution of new virulence traits.

### 3.4 The mitochondrial genome of *H. pseudoalbidus*

In order to identify sequences potentially originating from the mitochondrial genome of *H. pseudoalbidus* we downloaded the 248 fully sequenced ascomycete mitochondrial genomes from Genbank and used these sequences as a BLAST database to screen the genomic contigs for potential mitochondrial origin. Fifty-seven contigs were identified with significant similarity to ascomycete mitochondrial sequences. Further examination of these 57 contigs showed that many contigs were identical but in reverse complement or extending by a few hundred base pairs. These contigs were collapsed to form a dataset of 45 contigs ranging in length from 109-14,731 bp and GC-contents ranging from 9.2-45.9% (Figure 8). Most of the contigs *>*5 kb fall into a GC content range of 30-40%, typical of AT-rich mitochondrial sequences. It may be that the AT rich repeat islands discussed above are mitochondrial in origin as the mitochondrial genome will be more prevalent in the sequence dataset this would explain the increase in abundance of those sequences

**Figure 8:**
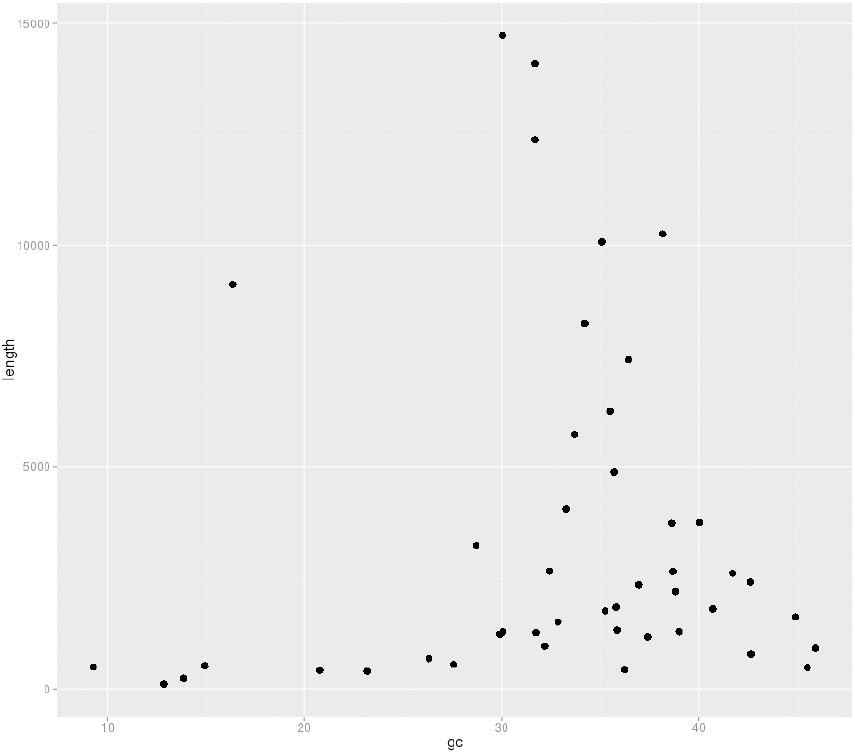
Contigs identified as potentially mitochondrial in origin by similarity searches we plot of length against GC content. The contigs with an unusually low GC content, <10%, appear to be composed of many AT rich repeats.Original available at http://dx.doi.org/10.6084/m9.figshare.1005003.

The total length of the 45 mitochondrial contigs is 156,026 bp with no significant overlap. If this preliminary estimate is accurate *H.pseudoalbidus* would have the largest mitochondrial genome sequenced from the ascomycetes so far (see Figure 9), although we expect the size to reduce with further work.

**Figure 9:**
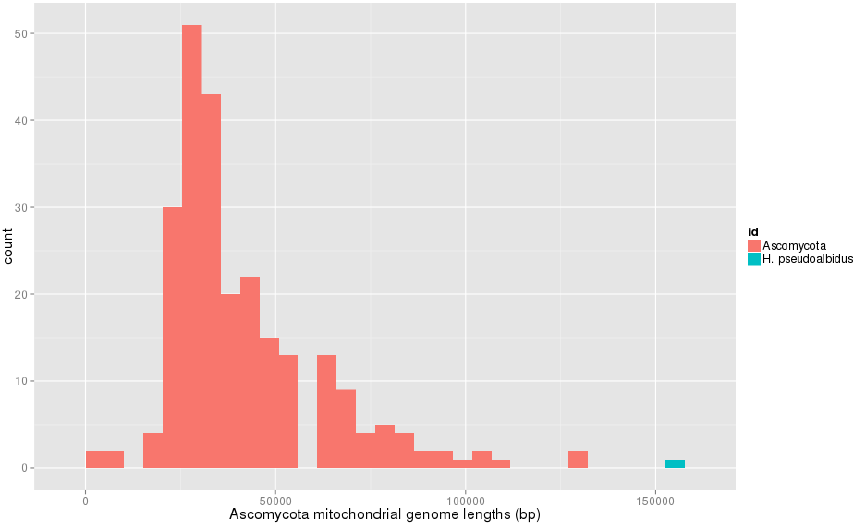
Histogram of mitochondrial genome lengths for all currently sequenced Ascomycete mitochondrial genomes. The combined length of the contigs currently identified as mitochondrial based on homology searches estimates the mitochondrial genome of *H. pseudoalbidus* at approximately 152 kb. Original available at http://dx.doi.org/10.6084/m9.figshare.1005004

A number of factors have prevented the construction of a finished mitochondrial genome at this time. Firstly, the potential mitochondrial contigs were identified based upon similarity based searches against current ascomycete mitochondrial genomes. The similarity based approach to finding mitochondrial sequences within a nuclear genome sequencing project may have misidentified some of these contigs as mitochondrial when in fact they are nuclear integrations of portions of the true mitochondrial genome (NUMTs). This is likely to have artificially inflated our estimate of the size of the *H. pseudoalbidus* mitochondrial genome. Annotation of the potential mitochondrial contigs is in progress and there are early indications of a very large number of introns (intronic ORFs) present in the mitochondrial genome of *H. pseudoalbidus*. The second complicating factor in attempting to assemble the mitochondrial genome at this time is the large number of AT repeats present in the sequences we have identified as being mitochondrial in origin. The repeats are likely to be collapsed and appear to be at the ends of the contigs we have identified, preventing further assembly without additional sequencing.

## 4 Diversity and Evolution of *Hymemoscyphus pseudoalbidus*

### 4.1 The repeat islands do not contain an over-representation of genes under positive selection pressure

In filamentous plant pathogens such as the late blight oomycete pathogen *Phytophthora infestans*, a repeat-driven expansion has created repeat and transposable element (TE) rich, gene-sparse regions that are distinct from the gene-dense conserved regions, known as a two-speed genome architecture. In *P. infestans*, repeat-rich regions are enriched with genes associated with virulence and are under positive selection as indicated by observed elevated *dN/dS* ratios [9, 8]. Determining the distance of a gene to its closest coding gene neighbours, (designated flanking intergenic regions, FIRs), can be used to determine whether a gene resides in a gene-dense or gene-sparse environment. Given that genes associated with pathogenicity tend to have long FIRs in pathogen genomes, genome architecture could be used to identify new candidate pathogenicity genes.

To investigate whether a similar organisation occurs in the genome of *Hymenoscyphus pseudoalbidus* we firstly identified candidate effector genes in the gene annotations. First we searched the predicted proteome of *H.pseudoalbidus* KW1 for potential secreted proteins using SignalP 2 [10] with parameters described in [11]. Proteins containing transmembrane domains and proteins with mitochondrial signal peptides were removed using TMHMM and TargetP, respectively. We then clustered all proteins based on sequence similarity using OrthoMCL [12]. We identified clusters of proteins (known as tribes) that contained at least one secreted protein. These 593 tribes were then used for all further analysis. Next, we annotated the protein tribes for known effector features as described in http://oadb.tsl.ac.uk/?p=622. Finally, we assigned an e-value to each feature within a tribe in order to rank tribes based on their likelihood of containing effector proteins [13]. The features associated with the top 100 ranked tribes are displayed in Figure 10.

**Figure 10:**
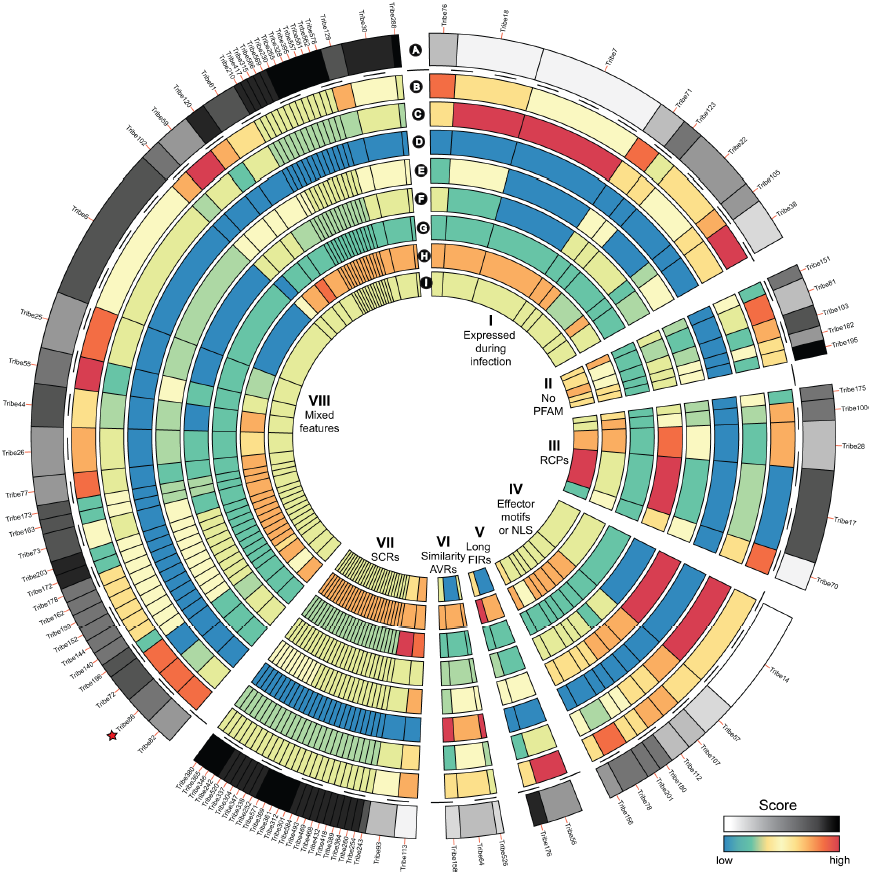
The top 100 ranked protein tribes containing putative effectors. A. Combined score used to rank tribes based on their content of effector features. B. Score for number of members classified as secreted. C. Score for number of members identified as expressed during infection. D. Score for number of members with similarity to known fungal AVRs. E. Score for number of members with effector motifs or nuclear localisation signals (NLS). F. Score for number of members classified as repeat containing. G. Score for number of members classified as small and cysteine rich. Score for number of members encoded by genes with at least one flanking intergenic region *>*10Kb. I. Score for number of members not annotated by PFAM domain searches. Red star indicates the tribe that contains potential Nep1-like protein (http://oadb.tsl.ac.uk/?p=235). Original available at http://dx.doi.org/10.6084/m9.figshare.1005005

In order to determine whether genes encoding secreted proteins are in gene sparse or dense regions of the genome we modified the *de novo* gene calls described in section 10.3 using RNA-seq data to extend based on overlaps with transcripts. Furthermore, the FIR distribution for genes in the *H.pseudoalbidus* genome can be seen in the Figure 11 and is indicative of a single speed genome, with genes encoding secreted proteins dispersed both in gene-sparse and gene-dense regions of the genome.

**Figure 11:**
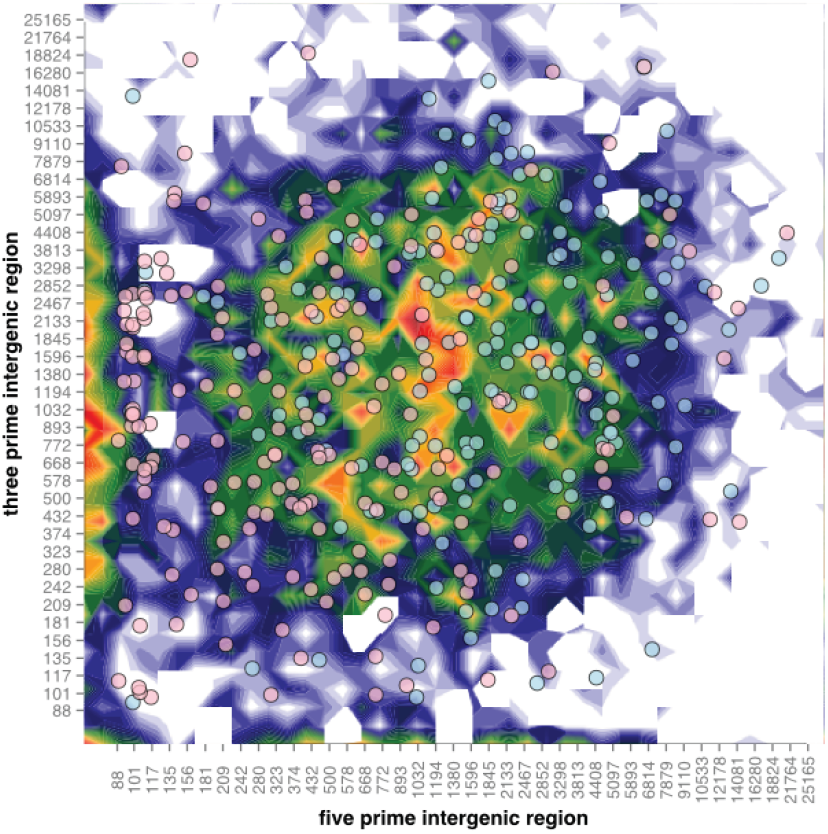
The single speed *H.pseudoalbidus* genome. Distribution of *H.pseudoalbidus* genes according to the length of their 5’ and 3’ flanking intergenic regions (FIRs). Red circles, core genes; blue circles, genes encoding predicted secreted proteins. Original available at http://dx.doi.org/10.6084/m9.figshare.1005006

As the AT-rich repeat regions do not appear to be assembled fully in the current draft assembly, we were not directly able to confirm whether sparseness of genes in these regions is an assembly artefact or feature of the genome and could not accurately assess genic content. As a proxy for this we examined the presence of small markers that represent genes, specifically considering the abundance of sequences corresponding to membrane localisation signal peptides or a nuclear localisation signal peptides. To identify whether the apparently collapsed repeats in the *H. pseudoalbidus* genome assembly may contain more secreted proteins and therefore be harbouring an enhanced number of effectors we searched the genome assembly for nucleotide sequences that may code for secretion or nuclear localisation signals and correlated the ratio of counts in 10 kbp sliding windows with the GC content of the same region in the genome assembly. We did not see a correlation between ratio of secretion signals to nuclear localisation signal and GC content of the and conclude that there is no enrichment of signal peptides, suggesting that the AT-rich regions of the genome are not enriched in secreted proteins and unlikely to harbour extra effectors (see supplemental figure at https://github.com/danmaclean/h_pseu_analysis/tree/master/nls_ss_gc).

### 4.2 Diversity between Hymenoscyphus pseudoalbidus isolates

To address the diversity within the UK and Japanese populations of *Hymenoscyphus pseudoalbidus*, we constructed a maximum likelihood phylogenetic tree of UK and Japanese isolates based on the third codon positions of all identified single copy genes. The tree in Figure 12 illustrates that Japanese and UK *H. pseudoalbidus* are clearly separated. This observation is consistent with previous studies based on several genes. To clarify genetic diversity in UK and Japanese *H. pseudoalbidus*, we estimated the nucleotide diversity (*π*) for coding regions of nuclear genes whose coverage of RNA sequencing short reads was over 90 percent (UK isolates: 5808 genes, Japanese isolates: 6140 genes). The average *π* for the tested genes is 0.0020 *±* 0.0091 for the UK isolates and 0.0098 *±* 0.0066 for the Japanese isolates, respectively. The Japanese isolates had higher nucleotide diversity compared with the UK isolates. We also estimated Tajima’s D to evaluate neutrality and population equilibrium in the UK *H. pseudoalbidus*. The average Tajima’s D value is −0.328 for UK isolates. The UK isolates showed 76 genes with significantly negative values (*p <*= 0.001), suggesting that these genes were under purifying selection or positive selection. Tajima’s D for 58 genes of the UK isolates was significantly positive (*p <*= 0.001). This positive value is expected under balancing selection. A McDonald and Kreitman (MK) test was performed to evaluate molecular evolution in the UK and Japanese isolates. In the neutral hypothesis of molecular evolution, the ratio of synonymous to non-synonymous divergences between the populations would be expected to be similar to the ratio of synonymous to non-synonymous polymorphisms within species. Since the tree showed clear separation between the UK and Japanese isolates, we used Japanese isolates as an out-group of the UK isolates. Only one gene encoding succinate dehydrogenase iron sulfur protein gave a statistically significant value (*p* = 0.048). Twenty synonymous differences within Japanese isolates were detected in this gene, while no polymorphism in the UK isolates. There was one non-synonymous difference between the UK and Japanese isolates, which is involved in a minor effect change from isoleucine to valine. The unusual number of synonymous polymorphisms in Japanese isolates caused this significance. The result of the McDonald Kreitman test suggests that the tested genes are unlikely to have been subjected to positive selection through the separation of the UK and Japanese isolates.

**Figure 12:**
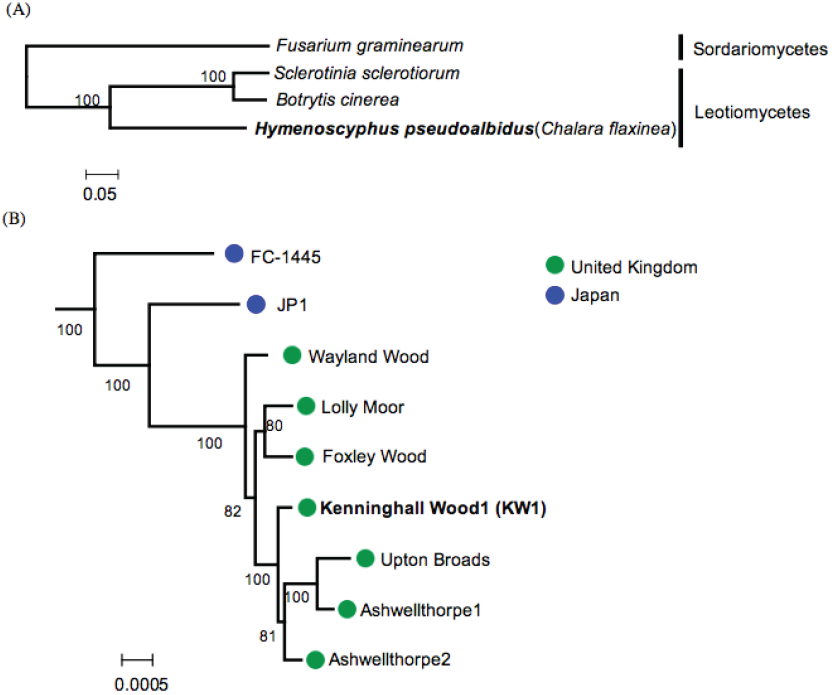
Maximum Likelihood trees for *H. pseudoalbidus* based on 3rd codon positions of 2,964 single copy genes. (A) Phylogenetic relationship between *H. pseudoalbidus* and fungi belonging to Sordariomycetes and Leotiomycetes. The subtree of *H. pseudoalbidus* was compressed. (B) Phylogenetic relationship between UK isolates and Japanese isolates. The number on the branches shows bootstrap support from 200 iterations. Original available at http://dx.doi.org/10.6084/m9.figshare.1005007

## 5 Metagenomic analysis of infected branch material

To check the species composition of samples from ash branch material, we performed a metagenomic analysis on several datasets that had been generated from samples described as *Fraxinus excelsior*, *H.pseudoalbidus* or as mixed infected material. We identified many transcripts specific to the infected samples (not in uninfected *Fraxinus*) that could not be assigned to the *H.pseudoalbidus* and *Fraxinus*; these may indicate additional microbial species present during infection perhaps acting synergistically with *H.pseudoalbidus* during infection or opportunists present as a secondary consequence of infection. We used the uninfected *F. excelsior* sample to identify species that are part of the ‘normal’ or ‘healthy’ tree microbiota and could therefore be excluded from the list of infection-related species. For each of the six samples, we identified the numbers of transcripts assigned to Helotiales and to Viridiplantae as described in section 10.10. The results are summarised in Table 3. Transcripts from the *H. pseudoalbidus* isolate KW1, assigned to Dothideomycetes, Eurotiomycetes, Leotiomycetes and Sordariomycetes, which all reside within the subphylum of Pezizomycotina, consistent with the sequenced sample being pure *Chalara*. As expected, all of the identifiable transcripts from the *F. excelsior* ATU1 transcripts fell within the Viridiplantae kingdom, specifically within the group of flowering plants (Magnoliophyta).

**Table 3:**
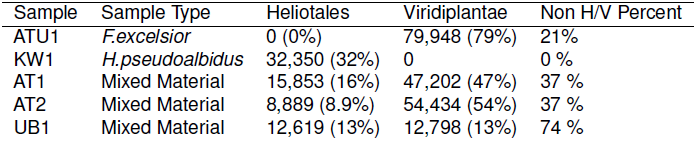
Reads assigned to Helotiales and Viridiplantae in normalised comparison for each sample examined. Total reads and percent of sample are shown. Full text version available at http://dx.doi.org/10.6084/m9.figshare.1005038

**Table 4:** Read number pre- and post-filtering, G + C content of raw data and percentage of KW1 genome positions covered at a depth of *>*10X.Full text version available at http://dx.doi.org/10.6084/m9.figshare.1005039

| Sample | Sample type | Raw reads x2 | GC% | Filtered x2 | % 10X KW1 genome positions covered |
| --- | --- | --- | --- | --- | --- |
| KW1 | *H. pseudoalbidus* | 41036951 | 49 | 38594279 | 32% |
| FERA 105 | *H. pseudoalbidus* | 1442015 | 48 | 1440015 | 0.26% |
| FERA 88 | *H. pseudoalbidus* | 173461 | 47 | 1729869 | 0.20% |
| FERA 93 | *H. pseudoalbidus* | 1355421 | 47 | 1353849 | 0.24% |
| FERA 94 | *H. pseudoalbidus* | 1642453 | 47 | 1639096 | 0.17% |
| FERA 232 | *H. pseudoalbidus* | 1919713 | 48 | 1918416 | 0.27% |
| FERA 233 | *H. pseudoalbidus* | 1838409 | 47 | 1837097 | 0.26% |
| Mature fruiting body (MFB1) | *H. pseudoalbidus* | 15033876 | 49 | 14524487 | 27% |
| Primordial fruiting body (PFB1) | *H. pseudoalbidus* | 7298316 | 49 | 7187891 | 16% |
| AT1 | Mixed material | 31059879 | 48 | 30388033 | 29% |
| AT2 | Mixed material | 23096021 | 44 | 21924824 | 13% |
| Holt | Mixed material | 36960651 | 49 | 36231834 | 21% |
| Upton | Mixed material | 43099662 | 48 | 41367071 | 32% |

**Table 5:** VarScan SNP calling of *H. pseudoalbidus* isolates, fruiting body and mixed material samples. Full text version available at http://dx.doi.org/10.6084/m9.figshare.1005040

| Sample | Sample type | Number Unambiguous bases | Number SNPs | VarScan Total |
| --- | --- | --- | --- | --- |
| KW1 | *H. pseudoalbidus* | 19974663 | 59 | NA |
| FERA 88 | *H. pseudoalbidus* | 167378 | 40 | 204 |
| FERA 93 | *H. pseudoalbidus* | 127827 | 20 |  |
| FERA 94 | *H. pseudoalbidus* | 150981 | 47 |  |
| FERA 105 | *H. pseudoalbidus* | 104667 | 22 |  |
| FERA 232 | *H. pseudoalbidus* | 169295 | 53 |  |
| FERA 233 | *H. pseudoalbidus* | 165989 | 35 |  |
| MFB1 | *H. pseudoalbidus* | 16519209 | 11852 | 16396 |
| PFB1 | *H. pseudoalbidus* | 9869668 | 9545 |  |
| AT12 | Mixed material | 18108777 | 6029 | NA |
| AT22 | Mixed material | 8016701 | 3138 |  |
| Holt | Mixed material | 12975549 | 13654 |  |
| Upton | Mixed material | 19789944 | 12259 |  |

**Table 6:** Assembly annotation statistics for different feature types. Full text version available at http://dx.doi.org/10.6084/m9.figshare.1005034

|  | Gene | mRNA | Exon | Intron | CDS | 5UTR | 3UTR | cDNA Transcript | CDS Transcript |
| --- | --- | --- | --- | --- | --- | --- | --- | --- | --- |
| Count | 10252 | 10405 | 35288 | 22795 | 33086 | 11954 | 10758 | 10405 | 10405 |
| Mean size | 2252 | 2255 | 590 | 83 | 478 | 201 | 242 | 2001 | 1520 |
| Minimum | 285 | 285 | 6 | 6 | 3 | 1 | 1 | 204 | 84 |
| Maximum | 25565 | 25565 | 14475 | 1305 | 14316 | 1798 | 1999 | 23327 | 22749 |
| Total Length | 23087543 | 23462930 | 20815617 | 1887836 | 15814549 | 2402584 | 2598484 | 20815617 | 15814549 |

In the data from Upton mixed material, 74% of the transcripts were not assigned to the Helotiales or Viridiplantae 13, which is where *H.pseudoalbidus*and *F. excelsior* transcripts are expected to fall, based on the results from pure isolate of *H.pseudoalbidus* and the uninfected *F. excelsior*. In the UB1 data, 34% of the total number of transcripts was assigned to Oomycetes; specifically 33% to *Phytophthora* spp.. Additionally, 13% were not assigned to any taxon and a further 11% had no significant similarity to proteins in the GenBank database, detectable by BLAST. The HC1 mixed material sample analysis showed 1.5% transcripts binned to *Togninia minima*, an ascomycete in the order Calosphaeriales. *T. minima* is a pathogen of grapevines and *Prunus* spp., however, the closely related *T. fraxinopennsylvanica* (anamorph: *Phaeoacremonium mortoniae*) has been observed in dead vascular tissue of declining ash tree branches (*Fraxinus latifolia*) in California [14, 15]. It may be that *T. fraxinopennsylvanica* is present in the HC1 material and that the reads were assigned to the species *T. minima* because that is the most closely related species for which extensive sequence data is available. For the nominally pure sample of *H.pseudoalbidus* isolate KW1, only 32% of reads were assigned to Helotiales. However, 66% of reads were assigned to Fungi with 16% not assigned to taxa and 17% had no significant similarity to proteins in the GenBank BLAST database. This is likely to be due to insufficient sequence data in the GenBank database from *H.pseudoalbidus* and closely related species.

## 6 Biology of Hymenoscyphus pseudoalbidus

### 6.1 Two mating types exist in the UK population of *Hymenoscyphus pseudoalbidus*

We used the KW1 genome described above and collated the partial sequences of SLA2 and APN2 proteins that likely flank the mating locus and the MAT1-1-3 and MAT1-2-1 proteins that have been identified in *Hymenoscyphus pseudoalbicans* and used these in a tblastn against the KW1 assembled contigs (with default settings) [16] The tblastn search highlighted a single contig that contained the SLA2, APN2 and MAT1-1-3 genes, suggesting the KW1 *H.pseudoalbidus* isolate is of the MAT1-1 mating type. The sequence between the SLA2 and MAT1-1-3 genes was then extracted and used in a blastx search against the NCBI nr database. This analysis identified a region with similarity to a DNA polymerase zeta catalytic subunit and enabled the first complete characterization of the MAT1-1-3 mating locus for *H.pseudoalbidus*.

We next took the partial sequences of MAT1-1-3 and MAT1-2-1 proteins from *H. pseudoalbidus* and used these in a second tblastn search against the AT1 and AT2 assembled transcripts. The tblastn search highlighted transcripts from both assemblies with similarity to the MAT1-2-1 gene and not MAT1-1-3. This suggests that the isolates AT1 and AT2 from Ashwellthorpe Lower Wood are of the MAT1-2 mating type and are therefore different to the KW1 *H.pseudoalbidus* isolate from Kenninghall Wood, which is of mating type MAT1-1. We confirmed this bioinformatic analysis with the PCR method in [16] on 24 samples of potentially infected ash trees from Ashwellthorpe lower wood and identified *H. pseudoalbidus* MAT 1-1 (MAT1-1-3) and MAT 1-2 (MAT-1-2-1) mating types from these samples. The size of the PCR products (see Figure 14) indicated that five samples were of the MAT1-1 mating type and three were of the MAT1-2 mating type showing that Ashwellthorpe wood alone contains both mating types. These results indicate the potential for sexual recombination to occur with the UK *H.pseudoalbidus* population.

**Figure 13:**
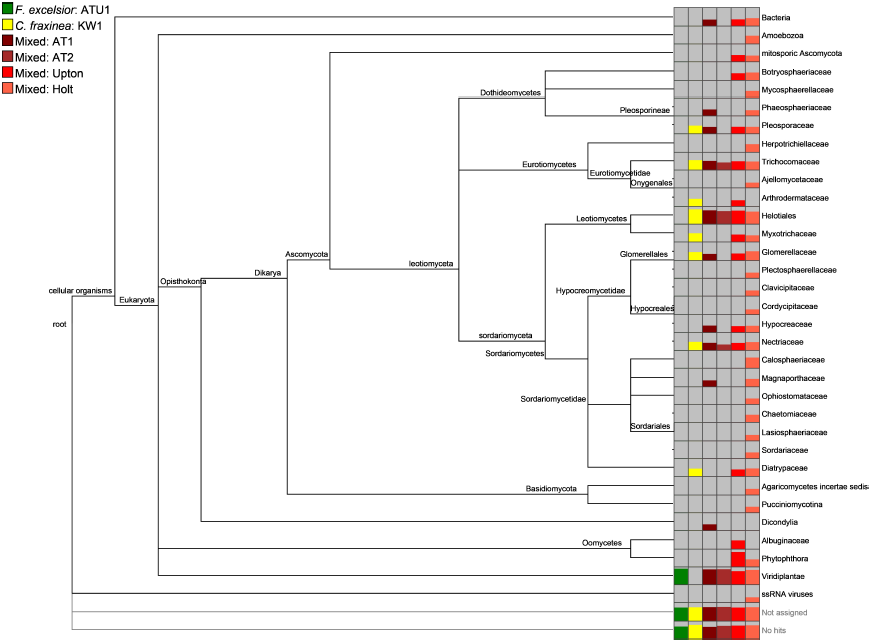
Metagenomic analysis of *F.excelsior*, *H.pseudoalbidus* and four infected material samples (AT1, AT2, UB1 and HP1). Original available at http://dx.doi.org/10.6084/m9.figshare.807684

**Figure 14:**
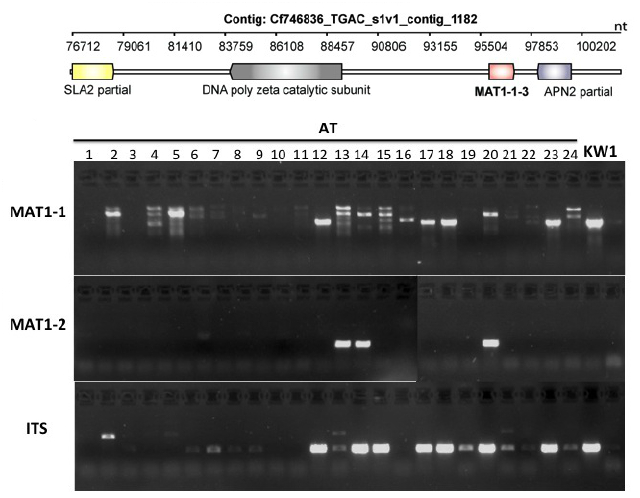
Mating-type locus analysis. Upper portion: The MAT locus in the KW1 isolate was identified in the genome sequence. A DNA polymerase zeta catalytic subunit was identified by tblastn of the region between the SLA2 and MAT1-1-3 genes against the NCBI nr database. Lower portion: *H.pseudoalbidus* PCR amplification of *H. pseudoalbidus* mating loci. AT indicates samples taken from Ashwellthorpe lower wood, KW1 is an isolate from Kenninghall wood that was previously confirmed as displaying the MAT1-1 (MAT1-1-3) mating type. ITS (internal transcribed spacer) analysis was used to confirm infection by *H.pseudoalbidus*. Original available at http://dx.doi.org/10.6084/m9.figshare.1005008

### 6.2 The genome of *Hymenoscyphus pseudoalbidus* does not contain genes for wood degradation like those seen in wood-rotting fungi

The infection process of *H.pseudoalbidus* remains unclear, infection may proceed to the pith by a number of potential routes including structures such as lenticels. To examine whether *H.pseudoalbidus* contains the apparatus to be able to metabolise woody tissue we compared the protein sequences with those of 31 other fungi including known wood-degrading basidiomycetes originally compiled in [17]. We used the list of proteins in the KW 1 genome assembly annotated in section (10.3 and a list of 38 wood decay related oxidoreductase and CAZyme [18] proteins derived by [17] as input to BLAST searches to identify proteins with strong sequence identity (see methods section 10.13). For each group in this decay-related list proteins were assayed to identify proteins with *>*50% sequence identity to the representative protein in at least 10 different species. Counts of the number of *H.pseudoalbidus* proteins passing this threshold were used as the estimate of the number of members of each group in the *H.pseudoalbidus* genome. The genome of *H.pseudoalbidus* does not appear to contain many proteins involved in wood degradation (see Figure 15) suggesting that it does not need to break down woody tissues to progress into the tree.

**Figure 15:**
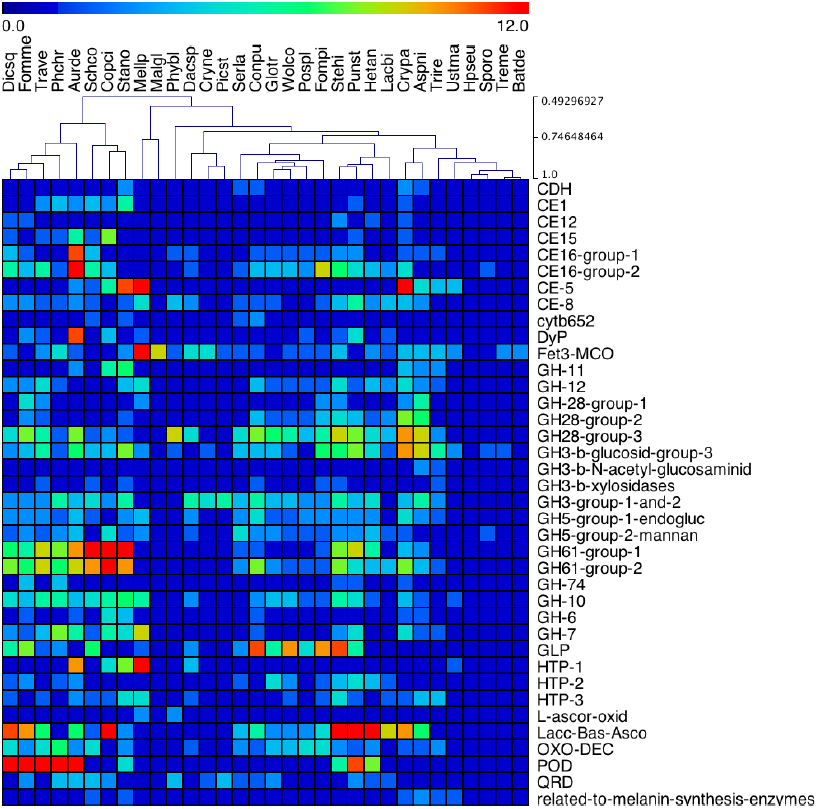
Heatmap showing counts of genes 11 oxidoreductase and 17 CAZyme [40] families in the genomes of 20 Agaricomycotina and 11 other fungi identified and counted in [17] extended to include *H.pseudoalbidus* - Hpseu, Hymenoscyphus pseudoalbidus; Aurde, Auricularia delicata; Conpu, Coniophora puteana; Dacsp, Dacryopinax sp., Dicsqs, Dichomitus squalens; Fomme, Fomitiporia mediterranea; Fompi, Fomitopsis pinicola; Gt, Gloeophyllum trabeum; Punst, Punctularia strigosozonata; Stehi, Stereum hirsutum; Treme, Tremella mesenterica, Trave, Trametes versicolor; Wolco, Wolfiporia cocos. Others: Aspni, Aspergillus niger; Batde, Batrachochytrium dendrobatidis; Copci, Coprinopsis cinerea; Cn, Cryptococcus neoformans; Crypa, Cryphonectria parasitica; Hetan, Heterobasidion annosum; Lacbi, Laccaria bicolor; Malgi, Malassezia globosa; Mellp, Melampsora laricis-populina; Phybl, Phycomyces blakesleeanus; Phchr, Phanerochaete chrysosporium; Pospl, Postia placenta; Picst, Pichia stipitis; Schco, Schizophyllum commune; Serls, Serpula lacrymans; Stano, Stagonospora nodorum; Sporo, Sporobolomyces roseus; Trire, Trichoderma reesei; Ustma, Ustilago maydis. Genes: GH, glycoside hydrolases; CE, carbohydrate esterases; POD, class II peroxidases; MCO, multicopper oxidases; CRO, copper-radical oxidases; CDH, cellobiose dehydrogenase; cytb562, cytochrome b562; OXO, oxalate oxidase/decarboxylases; GLP, Fe(III)-reducing glycopeptides; QRD, quinone reductases; DyP, dye-decolorizing peroxidases; HTP, heme-thiolate peroxidases. Original available at http://dx.doi.org/10.6084/m9.figshare.791635

### 6.3 The secretome of Hymenoscyphus pseudoalbidus

Secretion signal carrying proteins were analysed to identify Gene Ontology (GO) terms enriched relative to the entire proteome by hypergeometric test with *p <*= 0.05. Most notably enriched GO terms in the secretome of *H.pseudoalbidus* belong to those describing catalytic activity (GO:0003824) based on hydrolysis of oligomers of sugar or amino acid molecules (Supplemental Table 2). Cell wall degrading enzymes such as cellulases (GO:0008810), polygalacturonases (GO:0004650), pectate lyases (GO:0030570), glucosidases (GO:0015926), glucanases (GO:0052861), alpha-galactosidases (GO:0004557), pectinesterase (GO: 0030599), cutinases (GO:0050525), and alpha-L-fucosidase (GO:0004560) may be used by the pathogen to attack plant defenses and may be required for nutrient acquisition. Nutrients released by these enzymes could be captured by the proteins involved in nutrient reservoir activity (GO:0045735). These enzymes are necessarily secreted and may need to be maintained throughout the pathogen life cycle. *H.pseudoalbidus* also needs to maintain its own integrity by constantly repairing and recycling its cell wall perhaps using enzymes such as UDP-N-acetylmuramate dehydrogenase (GO:0008762) and chitinases (GO:0004568). As an invasive pathogen, *H.pseudoalbidus* may also degrade the plant proteins for source of amino acid and nitrogen for its development or it may need to neutralize plant proteins that are harmful for its progression. The fungus likely achieves this by secreting an array of peptidases (GO:0008233). The most enriched peptidases are serine peptidases (GO:0004252, GO:0008236) followed by aspartic endopeptidases (GO:0004190). Other types of peptidases such as Cysteine or metallo peptidases do not seem to be enriched in the secretome. The fungus also invests proteins with inhibitor activity (GO:0043086) in its secretomes presumably to control its own enzymes or to inhibit plant protein activity.

### 6.4 A Nep1-like protein (NLP) toxin in the *Hymenoscyphus pseudoalbidus* genome

We collated seven fungal and oomycete Nep1-like protein (NLP) sequences from public databases and used these in a tblastn against the AT1 assembled transcripts (with default settings). We identified one AT1 transcript with high similarity to NLPs, which contains all conserved residues and likely encodes a full length protein sequence. For this transcript we extracted the coding sequence and the corresponding protein sequence. The protein sequence was then used for protein structure modelling using Phyre2 [19], with the published NLPpya 3GNU [20] as a template. This analysis shows that the predicted protein we identified carries all the key residues for NLP cytolytic function as seen in Figure 16.

**Figure 16:**
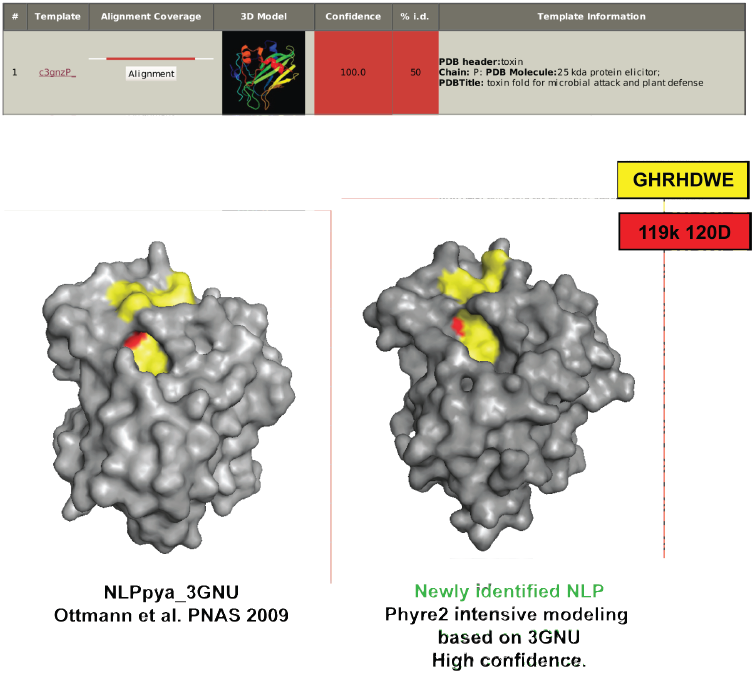
Protein structure modelling of sequence with identity to amino acid sequence of NLP-pya 3GNU [20].The known NLP is on the left, the model of the putative NLP is on the right. Original available at http://dx.doi.org/10.6084/m9.figshare.1005010

## 7 Key Achievements to Date

Our attempt to create a community response to ash dieback has been very fruitful. By releasing data early to permit analysis donated through crowdsourcing we have created a surprising amount of new knowledge in a relatively small amount of time. Our geographically disparate group collaborating primarily through OADB and using modern collaborative tools like GitHub has achieved a number of landmarks. Notably we have succeeded most in enabling genomic analysis of the causative agent of ash dieback *Hymenoscyphus pseudoalbidus*. The major achievements and findings of our crowdsourcing are:

1. Genomic DNA sequence and assembly of the *H.pseudoalbidus* genome, from an isolated culture originally taken from Kenninghall Wood in Norfolk, UK
2. Twelve assembled transcriptomes of *H.pseudoalbidus* from the UK and two from Japan
3. Annotations of the pathogen genome
4. Assessment of genetic diversity of the *H.pseudoalbidus* population in the UK

Through crowdsourcing we created a useable and informative ‘first-pass’ genome assembly with an excellent reconstruction of the gene space and a strong indication that the structure of the genome is interspersed with difficult to resolve AT-rich repeat islands. An interesting and unanswered question in our analyses concerns the structure of the mitochondrial genome, at least some of these repetitive contigs have strong identity to sequenced mitochondria and may belong in that genome. The current size estimate is likely to be inflated and we expect that finishing and improvement of the sequence we have will give us a clearer picture.

By making use of the crowdsourced annotations on the genome we have seen that the canonical disease causing proteins, effectors, do not appear to be encoded by genes in particular regions of, and have been able to rule out the presence of a ‘two-speed’ genome for *H.pseudoalbidus*. Furthermore, the quickly and freely provided annotations have provided domain experts the opportunity to search for toxin pathways. The phylogenetic analyses carried out on samples of *H.pseudoalbidus* from the UK and Japan suggest that the UK isolates represent a discrete clade, distinct from, but likely derived distantly from relatives of the Japanese isolates. Analysis of the ratio of nucleotide substitutions in the UK clade does not show evidence that it is undergoing any particular directional selection.

Together these analyses represent a significant success for crowdsourcing as a method of bringing a new community into existence, the approach outlined here has indeed brought the data to the people who can use it quickest [21] and swiftly delivered new information.

## 8 A future for crowdsourcing in genomics

Crowdsourcing as a bottom-up activity gains its strength from the essentially charitable donations of time and materials from interested parties. In contrast to crowdsourcing we can define a term ‘grantsourcing’ for the financially front-loaded way of achieving research that is most common across science. We have seen that the granular nature of the contributions means that extremely time-intensive or costly work is much less likely to be achieved through crowdsourcing alone. For example, the finishing of the genome sequence of *H.pseudoalbidus* requires effort and monetary investment greater than what will reasonably be achieved through crowdsourcing but is within the bounds that we can expect to be achieved from grantsourced science.

We expect that a marriage between crowdsourcing and grantsourcing will likely provide a speed/cost sweet-spot for the identification and execution of critical future research projects. The plant pathology community faces significant challenges as land use, trading practice, agricultural practice, climate and human population levels all change. Many more pathogen emergences are expected and a purely grantsourced approach is going to provide too little too late. There is definitely a place for crowdsourcing in the front line of pathogenomics. Crowdsourcing helps us to get projects started as soon as possible. Open science approaches [22] will help us maintain maximum speed as these studies progress and as we are able to show promising lines of further study, generate collaborative networks and align research interests we can more quickly and affirmatively move onto grantsource appropriate projects. It is clear to us that the key for maximising the speed at which we can carry out necessary research as a community lies in embracing and maintaining Open Science approaches to keep data and ideas flowing regardless of the source of funding or manpower by which the work gets done. The results of our crowdsourcing initiative shows that working in this way is feasible and productive.

## 9 Acknowledgements

We wish to thank Ghanasyam Rallapalli for technical assistance. We also and especially wish to thank all contributors to the OADB GitHub repository whose work is not covered in this report or who declined to be included as co-author from modesty about the size and importance of their help: all contributions are valuable and all are equally appreciated. The work in this document and on OADB has been funded by the Biotechnology and Biological Sciences Research Council and the Department for Environment, Food and Rural Affairs through the NORNEX project, The Gatsby Charitable Foundation, the John Innes Foundation, The Genome Analysis Centre National Capability Grant, a John Innes Foundation Emeritus Fellowship to JAD and a Nuffield Science Bursary to DB.

## 10 Methods

### 10.1 Assembly of short read sequences

Illumina short reads were assembled into scaffolds using ABySS [23] version 1.3.4 with the following command-line:

~~~
abyss-pe \
c = 10 \
np = 200 \
k = 202 \
lib="pe1 pe2” \
pe1="$READS_PE” \
pe2="$READS_OPE”
~~~

The resulting assembly was deposited in http://oadb.tsl.ac.uk GitHub repository at https://github.com/ash-dieback-crowdsource/data/blob/master/ash_dieback/chalara_fraxinea/Kenninghall_wood_KW1/assemblies/gDNA/KW1_assembly_version1/Chalara_fraxinea_TGAC_s1v1_scaffolds.fa

### 10.2 Statistical analysis of assembly quality

NG(X) graphs describing the distribution of scaffold lengths were calculated as described in [5], namely:

~~~
[NG(X)] is calculated by summing all sequence lengths,
starting with the longest, and observing the length that
takes the sum length past [X] % of the total assembly length.
~~~

Ruby and R code and data to replicate this analysis are provided in the public GitHub repository: https://github.com/danmaclean/h_pseu_analysis/tree/master/scaffold_size.

### 10.3 Evaluation of assembly of gene space

CEGMA [6] was used to assess the completeness of the gene space assembled using default settings

~~~
cegma -g Chalara_fraxinea_TGAC_s1v1_scaffolds.fa
~~~

using the scaffold file available at https://github.com/ash-dieback-crowdsource/data/blob/master/ash_dieback/chalara_fraxinea/Kenninghall_wood_KW1/assemblies/gDNA/KW1_assembly_version1/Chalara_fraxinea_TGAC_s1v1_scaffolds.fa Output from this run can be seen in the GitHub repository at: https://github.com/danmaclean/h_pseu_analysis/tree/master/cegma

### 10.4 Repeat Masking for Annotation

RepeatModeler v1-0-7 [24] was used to generate a species specific repeat library based on the KW1 assembly. Interspersed repeats were identified using the KW1 repeat library and RepeatMasker [25]. Low complexity repeats were identified with RepeatMasker and Tantan [26]. GFF files describing the locations of these features were deposited in the http://oadb.tsl.ac.uk GitHub repository at https://github.com/ash-dieback-crowdsource/data/tree/master/ash_dieback/chalara_fraxinea/Kenninghall_wood_KW1/annotations/Repeats/Repeatmasker https://github.com/ash-dieback-crowdsource/data/tree/master/ash_dieback/chalara_fraxinea/Kenninghall_wood_KW1/annotations/Repeats/Tantan

### 10.5 Protein evidence based sequence annotation

Protein sequences from 4 species:

1. Amorphotheca resinae v1.0,
2. Botrytis cinerea v1.0,
3. Oidiodendron maius Zn v1.0,
4. Sclerotinia sclerotiorum v1.0

from http://genome.jgi.doe.gov/pages/dynamicOrganismDownload.jsf?organism=leotiomycetes were soft masked for low complexity using segmasker from the BLAST+ package [27] and aligned to the soft-masked KW1 assembly with exonerate [28] protein2genome, alignments were filtered at a minimum 50% identity and 50% coverage. GFF files describing the locations of these features were deposited in the http://oadb.tsl.ac.uk GitHub repository at

1. https://github.com/ash-dieback-crowdsource/data/blob/master/ash_dieback/chalara_fraxinea/Kenninghall_wood_KW1/annotations/Sources/TGAC_Chalara_fraxinea_ass_s1v1_ann_v1.1/Amore1_GeneCatalog_proteins_20110719.aa.mask.exonerate.50id-50cov.gff
2. https://github.com/ash-dieback-crowdsource/data/blob/master/ash_dieback/chalara_fraxinea/Kenninghall_wood_KW1/annotations/Sources/TGAC_Chalara_fraxinea_ass_s1v1_ann_v1.1/Botci1_GeneCatalog_proteins_20110903.aa.mask.exonerate.50id-50cov.gff
3. https://github.com/ash-dieback-crowdsource/data/blob/master/ash_dieback/chalara_fraxinea/Kenninghall_wood_KW1/annotations/Sources/TGAC_Chalara_fraxinea_ass_s1v1_ann_v1.1/Oidma1_GeneCatalog_proteins_20110606.aa.mask.exonerate.50id-50cov.gff
4. https://github.com/ash-dieback-crowdsource/data/blob/master/ash_dieback/chalara_fraxinea/Kenninghall_wood_KW1/annotations/Sources/TGAC_Chalara_fraxinea_ass_s1v1_ann_v1.1/Sclsc1_GeneCatalog_proteins_20110903.aa.mask.exonerate.50id-50cov.gff

### 10.6 RNA-Seq based evidence annotation

The AT2 RNA-Seq reads were aligned to the unmasked TGAC KW1 assembly with Tophat version 2.0.4 [29] with the following options

~~~
--min-anchor-length 12
--max-multihits 20
--min-intron-length 50
--max-intron-length 10000
--min-coverage-intron 50
--max-coverage-intron 5000
--min-segment-intron 50
--max-segment-intron 10000
~~~

Splice junctions identified with Tophat [29] were filtered requiring support from a minimum of 4 reads. Junction file (unfiltered): https://github.com/ash-dieback-crowdsource/data/blob/master/ash_dieback/chalara_fraxinea/Kenninghall_wood_KW1/annotations/Sources/jasmbl_filtered.gbrowse.gff

A bigwig file of RNA-Seq read density is provided at: https://github.com/ash-dieback-crowdsource/data/blob/master/ash_dieback/chalara_fraxinea/Kenninghall_wood_KW1/annotations/Source/accepted_hits.sort.bam.update.bw

RNA-Seq alignments were assembled with Cufflinks version 2.0.2 [30] using options

-I 10000

-A 0.15

from this assembly the longest ORFs were selected: https://github.com/ash-dieback-crowdsource/data/blob/master/ash_dieback/chalara_fraxinea/Kenninghall_wood_KW1/annotations/Source/cufflinks_transcripts_cds.gff

A trinity assembly of the same RNA-Seq data was prepared using trinity version 5-19-2011 [31] and command-line

~~~
Trinity.pl --seqType fq \
--left lane4_NoIndex_R1.fastq \
--right lane4_NoIndex_R2.fastq \
--output lane4_RNAseq_Trinity \
--run_butterfly
~~~

The resulting assembly was deposited in http://oadb.tsl.ac.uk GitHub repository at https://github.com/ash-dieback-crowdsource/data/tree/master/ash_dieback/mixed_material/ashwellthorpe_AT2/assemblies

The trinity assembly was softmasked (using dustmasker [27]) and aligned to the softmasked KW1 assembly with exonerate [28] est2genome and filtered at a minimum of 95% identity, 50% coverage.

GFF files describing the locations of these features were deposited in the OADB GitHub repository at https://github.com/ash-dieback-crowdsource/data/blob/master/ash_dieback/chalara_fraxinea/Kenninghall_wood_KW1/annotations/Source/AT2_trinity_version2.mask.fasta.exonerate.95id-50cov.gff

### 10.7 Augustus Training and genebuild

Augustus version 2.5.5 [32] was trained using a subset of cufflinks assemblies. Briefly, blast searches against the 4 protein datasets in section 10.5, were used to identify predicted cufflinks CDS features with support from cross species alignments, these were further filtered to remove features showing over 80% sequence similarity within the selected subset or any genomic overlap. Augustus was trained based on the filtered set of 859 cufflinks models with 100 models reserved for testing. Based on this the *ab initio* predictions achieved sensitivity results of 0.96 nucleotide, 0.80 exon and 0.65 at the gene level.

Augustus gene models were predicted using the trained *ab initio* model with the 4 cross species protein alignments (section 10.5), RNA-Seq junctions (section 10.6), Cufflinks alignments (section 10.6), and aligned trinity contigs (section 10.6) as evidence hints. RNA-Seq read density was provided as exon hints and repeat information as nonexonpart hints.Augustus gene models were filtered to remove redundant genes giving a total of 10252 predicted genes with 10405 predicted mRNAs. These final gene models were assigned a unique identifier of the following form CHAFR746836.1.1_0000010.1, (NCBI taxonomy id: CHAFR746836, annotation version: .1.1, gene identifier: _0000010, transcript identifier:.1).

GFF files describing the locations of these features were deposited in the OADB GitHub repository at https://github.com/ash-dieback-crowdsource/data/blob/master/ash_dieback/chalara_fraxinea/Kenninghall_wood_KW1/annotations/Gene_predictions/TGAC_Chalara_fraxinea_ass_s1v1_ann_v1.1/Chalara_fraxinea_ass_s1v1_ann_v1.1.gene.gff

### 10.8 Gene Ontology annotation of protein sequences

Protein sequences were analysed using the EBI webservice InterproScan and the EBI provided Perl script iprscan5_lwp.pl. Perl, Ruby and R code and data to replicate this analysis and supplemen-tal results are provided in the public GitHub repository at https://github.com/danmaclean/h_pseu_analysis/tree/master/go_analysis

### 10.9 Alignment of gDNA reads to KW1 reference

The genomic DNA sequence reads referenced in section 10.18 were aligned with bwa [33], and the scaffold assembly described in 10.1 using the bwa-mem [34] algorithm and default alignment settings at all steps.

### 10.10 Metagenome analysis

We used MEGAN [35] and the assembled transcripts from *F. excelsior* and *H.pseudoalbidus*to identify the taxonomic groups to which uninfected sample transcripts are allocated, to use as a reference database for binning.We identified sequence similarity between assembled transcripts and GenBank protein sequences using BLASTX; we used as queries the transcripts from uninfected *F. excelsior* (ATU1), a *H.pseudoalbidus* isolate (KW1) and three mixed material samples (AT1, AT2, Upton). We loaded the output from BLASTX into MEGAN and performed taxonomic binning using a minimum support value of 35, a minimum BLAST score of 50 and only retaining hits whose bit scores lie within 10% of the best score. The analyses were normalised, compared and rendered within MEGAN.

### 10.11 Preparation of Circos plots

Figures of KW1 assembly scaffolds were rendered in Circos version 0.64 [36]. Data, scripts and config files to replicate the figure generation is provided in the GitHub repository at https://github.com/danmaclean/h_pseu_analysis/blob/master/circos/scripts/assess_insert_size_distributions.md

### 10.12 Signal and Nuclear localisation Peptide identification and analysis

We extracted all open-reading frames from the KW1 assembly scaffold nucleotide sequence at https://github.com/ash-dieback-crowdsource/data/blob/master/ash_dieback/chalara_fraxinea/Kenninghall_wood_KW1/assemblies/gDNA/KW1_assembly_version1/Chalara_fraxinea_TGAC_s1v1_scaffolds.fa then searched the 6-frame translations of these for sequences predicted to code for secretion signals with SignalP 4 and independently for nuclear localisation signals with NLStradamus doi:10.1186/1471-2105-10-202, using a local Galaxy instance and custom scripts. R, Ruby code and data files and a Galaxy Workflow to replicate this analysis are provided in this GitHub repository https://github.com/danmaclean/h_pseu_analysis/tree/master/nls_ss_gc.

### 10.13 Identification of lignin-digestion related enzymes

The protein annotations from 10.5 were used in searches with BLAST+ version 2.2.8 [27] against sequence databases from [17]. For each group in this decay-related list *H.pseudoalbidus* proteins were assayed to identify proteins with *>*50% sequence identity to the representative protein in at least 10 different species. Counts of the number of *H.pseudoalbidus* proteins passing this threshold were used as the estimate of the number of members of each group in the *H.pseudoalbidus* genome. Code and data files to replicate this analysis are provided in the GitHub repository at https://github.com/danmaclean/h_pseu_analysis/tree/master/interesting_wood_genes

### 10.14 SNP identification from RNA-seq reads

RNA-seq reads were aligned to the genomic DNA assembly described in section 10.1 in tophat version 2.0.8 with the GFF of gene models described in section 10.3 as an input. The resulting alignment BAM files were then used in GATK version 2.7.4 to call SNPs and INDELS. Ruby scripts and command-lines used to reproduce this analysis are available at https://github.com/danmaclean/h_pseu_analysis/tree/master/tophat_gatk.

### 10.15 Construction of phylogenetic trees

We reconstructed consensus sequences of coding regions based on alignments of RNA-seq reads. Heterozygous sites are expressed using IUPAC (International Union of Pure and Applied Chemistry) codes. We only used 3rd codon positions that constituted 80% coverage of the tested samples for the construction of phylogenetic trees. The Maximum likelihood tree was constructed using RAxML software.

### 10.16 Estimation of nucleotide diversity and test of neutrality

We presumed haplotypes based on the consensus sequences. Each sample was virtually regarded as a single diploid isolate. After discarding genes whose unclear bases were over 20% of total bases of coding regions, 18 haplotypes (nine European samples) for 5182 cDNAs were predicted. We estimated nucleotide diversity *π*, performed Tajima’s D tests and McDonald and Kreitman tests using the software libsequence [37].

### 10.17 SNP identification for comparison of isolates and fruiting bodies

Qualities of raw reads from the thirteen samples were assessed with FASTQC [38]. Adapter- and quality-trimming was performed with Trim galore, a wrapper script using FASTQC and cutadapt (phred cutoff:20, error rate:0.1, adapter overlap: 1 bp, min. length: 20 bp, paired read length cut-off: 35 bp). FASTQC was automatically run on all trimmed files to confirm trimmed read quality. Raw and trimmed metrics and GC content of raw reads can be seen in Table 1.

Trimmed reads were aligned to the pre-assembled KW1 genome from OADB using the splice-aware aligner Tophat [29] The resulting assembly BAM file was used to create a pileup file using MPILEUP from SAMtools [33].

The variant detection software, VarScan2 [39] pileup2snp (or mpileup2snp) was used (–p-value 0.05–min-coverage 10–output-vcf-min-var-freq 0.95) to call SNPs. The numbers of SNPs were normalised to the number of bases with ¿=10X coverage to take into account the different depths of sequencing.

### 10.18 Genomic DNA sequence read accessions

KW1 Genomic DNA sequence read sets used in the KW1 genomic assembly are currently available by anonymous FTP from the following URLs:

1. ftp-oadb.tsl.ac.uk/chalara_fraxinea/Kenninghall_wood_KW1/reads/read_set_1/lib2569.fastq.tar.gz
2. ftp-oadb.tsl.ac.uk/chalara_fraxinea/Kenninghall_wood_KW1/reads/read_set_2/lib2570.fastq.tar.gz

### 10.19 RNA sequence read accessions

AT2 RNA-seq reads from mixed material used in evidence based transcript annotation in 10.6 and SNP and INDEL calling 10.14 Collected at geographical location: ‘52.538387,1.152577’

1. ftp-oadb.tsl.ac.uk/mixed_material/ashwellthorpe_AT2/RNA_seq/lane5_NoIndex_L005_R1_001.fastq.gz
2. ftp-oadb.tsl.ac.uk/mixed_material/ashwellthorpe_AT2/RNA_seq/lane5_NoIndex_L005_R1_002.fastq.gz
3. ftp-oadb.tsl.ac.uk/mixed_material/ashwellthorpe_AT2/RNA_seq/lane5_NoIndex_L005_R1_003.fastq.gz
4. ftp-oadb.tsl.ac.uk/mixed_material/ashwellthorpe_AT2/RNA_seq/lane5_NoIndex_L005_R1_004.fastq.gz
5. ftp-oadb.tsl.ac.uk/mixed_material/ashwellthorpe_AT2/RNA_seq/lane5_NoIndex_L005_R1_005.fastq.gz
6. ftp-oadb.tsl.ac.uk/mixed_material/ashwellthorpe_AT2/RNA_seq/lane5_NoIndex_L005_R1_006.fastq.gz
7. ftp-oadb.tsl.ac.uk/mixed_material/ashwellthorpe_AT2/RNA_seq/lane5_NoIndex_L005_R2_001.fastq.gz
8. ftp-oadb.tsl.ac.uk/mixed_material/ashwellthorpe_AT2/RNA_seq/lane5_NoIndex_L005_R2_002.fastq.gz
9. ftp-oadb.tsl.ac.uk/mixed_material/ashwellthorpe_AT2/RNA_seq/lane5_NoIndex_L005_R2_003.fastq.gz
10. ftp-oadb.tsl.ac.uk/mixed_material/ashwellthorpe_AT2/RNA_seq/lane5_NoIndex_L005_R2_004.fastq.gz
11. ftp-oadb.tsl.ac.uk/mixed_material/ashwellthorpe_AT2/RNA_seq/lane5_NoIndex_L005_R2_005.fastq.gz
12. ftp-oadb.tsl.ac.uk/mixed_material/ashwellthorpe_AT2/RNA_seq/lane5_NoIndex_L005_R2_006.fastq.gz

AT1 RNA-seq reads from mixed material used in SNP and INDEL analysis 10.14 Collected at geographical location: ‘52.538387,1.152577’

1. ftp-oadb.tsl.ac.uk/mixed_material/ashwellthorpe_AT1/RNA_seq/lane4_NoIndex_L004_R1_001.fastq.gz
2. ftp-oadb.tsl.ac.uk/mixed_material/ashwellthorpe_AT1/RNA_seq/lane4_NoIndex_L004_R1_002.fastq.gz
3. ftp-oadb.tsl.ac.uk/mixed_material/ashwellthorpe_AT1/RNA_seq/lane4_NoIndex_L004_R1_003.fastq.gz
4. ftp-oadb.tsl.ac.uk/mixed_material/ashwellthorpe_AT1/RNA_seq/lane4_NoIndex_L004_R1_004.fastq.gz
5. ftp-oadb.tsl.ac.uk/mixed_material/ashwellthorpe_AT1/RNA_seq/lane4_NoIndex_L004_R1_005.fastq.gz
6. ftp-oadb.tsl.ac.uk/mixed_material/ashwellthorpe_AT1/RNA_seq/lane4_NoIndex_L004_R1_006.fastq.gz
7. ftp-oadb.tsl.ac.uk/mixed_material/ashwellthorpe_AT1/RNA_seq/lane4_NoIndex_L004_R1_007.fastq.gz
8. ftp-oadb.tsl.ac.uk/mixed_material/ashwellthorpe_AT1/RNA_seq/lane4_NoIndex_L004_R1_008.fastq.gz
9. ftp-oadb.tsl.ac.uk/mixed_material/ashwellthorpe_AT1/RNA_seq/lane4_NoIndex_L004_R2_001.fastq.gz
10. ftp-oadb.tsl.ac.uk/mixed_material/ashwellthorpe_AT1/RNA_seq/lane4_NoIndex_L004_R2_002.fastq.gz
11. ftp-oadb.tsl.ac.uk/mixed_material/ashwellthorpe_AT1/RNA_seq/lane4_NoIndex_L004_R2_003.fastq.gz
12. ftp-oadb.tsl.ac.uk/mixed_material/ashwellthorpe_AT1/RNA_seq/lane4_NoIndex_L004_R2_004.fastq.gz
13. ftp-oadb.tsl.ac.uk/mixed_material/ashwellthorpe_AT1/RNA_seq/lane4_NoIndex_L004_R2_005.fastq.gz
14. ftp-oadb.tsl.ac.uk/mixed_material/ashwellthorpe_AT1/RNA_seq/lane4_NoIndex_L004_R2_006.fastq.gz
15. ftp-oadb.tsl.ac.uk/mixed_material/ashwellthorpe_AT1/RNA_seq/lane4_NoIndex_L004_R2_007.fastq.gz
16. ftp-oadb.tsl.ac.uk/mixed_material/ashwellthorpe_AT1/RNA_seq/lane4_NoIndex_L004_R2_008.fastq.gz

Upton Broad and Marshes RNA-seq reads from mixed material used in SNP and INDEL analysis 10.14 Collected at geographical location: ‘52.663253,1.533494’

1. ftp-oadb.tsl.ac.uk/mixed_material/Upton_broad_and_Marshes_UB1/reads/RNAseq/TSL_ID153_lane2/lane2_NoIndex_R1.fastq
2. ftp-oadb.tsl.ac.uk/mixed_material/Upton_broad_and_Marshes_UB1/reads/RNAseq/TSL_ID153_lane2/lane2_NoIndex_R2.fastq Lolly Moor RNA-seq reads from mixed material used in SNP and INDEL analysis 10.14 Collected at geographical location:‘52.613969,0.886402’
3. ftp-oadb.tsl.ac.uk/mixed_material/Lollymoor_LM1/lane6_ACAGTG_R1.fastq
4. ftp-oadb.tsl.ac.uk/mixed_material/Lollymoor_LM1/lane6_ACAGTG_R2.fastq

Japanese 1 RNA-seq reads from mixed material used in SNP and INDEL analysis 10.14

1. ftp-oadb.tsl.ac.uk/mixed_material/Japan/JP2/lane3_CAGATC_R1.fastq
2. ftp-oadb.tsl.ac.uk/mixed_material/Japan/JP2/lane3_CAGATC_R2.fastq

Japanese 2 RNA-seq reads from mixed material used in SNP and INDEL analysis 10.14

1. ftp-oadb.tsl.ac.uk/mixed_material/Japan_JP1/lane3_ACAGTG_R1.fastq
2. ftp-oadb.tsl.ac.uk/mixed_material/Japan_JP1/lane3_ACAGTG_R2.fastq

Holt Country Park RNA-seq reads from mixed material used in SNP and INDEL analysis 10.14 Collected at geographical location: ‘52.893888,1.095071’

1. ftp-oadb.tsl.ac.uk/mixed_material/Holt_Country_Park_HP1/reads/RNAseq/TSL_ID153_lane1/lane1_NoIndex_R1.fastq
2. ftp-oadb.tsl.ac.uk/mixed_material/Holt_Country_Park_HP1/reads/RNAseq/TSL_ID153_lane1/lane1_NoIndex_R2.fastq

Foxley Wood RNA-seq reads from mixed material used in SNP and INDEL analysis 10.14 Collected at geographical location:‘52.761092,1.042749’

1. ftp-oadb.tsl.ac.uk/mixed_material/Foxley/lane6_CAGATC_R1.fastq
2. ftp-oadb.tsl.ac.uk/mixed_material/Foxley/lane6_CAGATC_R2.fastq

