## Supplemental Table for "Crowdsourced analysis of ash and ash dieback through the Open Ash Dieback project: A year 1 report on datasets and analyses contributed by a self-organising community"

| Date | Committer | Commit Comment |
|---|---|---|
| 29/11/2013 | Shyam Rallapalli | Updated analysis markdown |
| 29/11/2013 | Shyam Rallapalli | update of PFAM domain and Gene Ontology for Chalara protein predictions |
| 25/11/2013 | Martin Trick | Repeated sample of Tree35 RNA-Seq reads together with assemblies annotation and BLAST results. Constitutes a replacement canonical reference sequence for use in association studies |
| 20/11/2013 | Shyam Rallapalli | blastn to gff info added gff file was generated from matrintrick analysis of blastall blastn of RNA-seq assembly |
| 20/11/2013 | Shyam Rallapalli | RNA-seq blastn to GFF annotations tree35 and ATU1 RNA-seq trinity assemblies are used for blast+ blastn and alignment locaitons are presented as gff |
| 08/11/2013 | Shyam Rallapalli | Go ids appended to GFF Inteproscan5 analysis used to append Go id information to GFF and SVG file prodcuded by the analysis are stored |
| 08/11/2013 | Shyam Rallapalli | Interproscan 5 of predicted proteins Interproscan 5 analysis created SVG files for predicted proteins Chalara. SVG files for the 5961 proteins are stored in SVG-out folder |
| 24/09/2013 | Dan MacLean | Variants for Foxley Wood Lolly Moor and Japanese samples Added GATK called SNPs and INDELS from Foxley Wood Lolly Moor and Japanese samples collected from mixed material |
| 24/09/2013 | Diane Saunders | Reads added for RNAseq data generated from a Japanese sample (sample 2) |
| 24/09/2013 | Diane Saunders | Reads added for RNAseq data generated from a Japanese sample (sample 1) |
| 24/09/2013 | Diane Saunders | Reads added for RNAseq data generated from a sample from Foxley Wood |
| 24/09/2013 | Diane Saunders | Reads added for RNAseq data generated from a sample from Lolly Moor. |
| 13/09/2013 | ethering | Moved GIRM2 and LSVM82 to h_pseudoalbidus |
| 10/09/2013 | Diane Saunders | Effector mining spreadsheet added |
| 28/08/2013 | rachelglover | Blastn of FERA_94 assembly vs. KW1 mycelia assembly |
| 28/08/2013 | rachelglover | Blastn of FERA_93 assembly vs. KW1 mycelia assembly |
| 28/08/2013 | rachelglover | Blastn of FERA_233 assembly vs. KW1 mycelia assembly |
| 28/08/2013 | rachelglover | Blastn of FERA_232 assembly vs. KW1 mycelia assembly |
| 28/08/2013 | rachelglover | Blastn of FERA_105 assembly vs. KW1 mycelia assembly |
| 28/08/2013 | rachelglover | Blastn of FERA_88 assembly against Kenninghall wood mycelia assembly |
| 28/08/2013 | Rachel Glover | Identified mitovirus sequences from their respective assemblies |
| 28/08/2013 | Rachel Glover | Blastx of AT1 assembly against genbank |
| 28/08/2013 | Rachel Glover | Blastx of AT2 assembly against genbank |
| 28/08/2013 | Rachel Glover | Blastx of AT1 assembly against Genbank |
| 28/08/2013 | Rachel Glover | Blastx of Upton assembly against genbank |
| 28/08/2013 | Rachel Glover | Blastx of Holt country park assembly against Genbank |
| 28/08/2013 | Rachel Glover | Blastx of Ashwellthorpe ATU1 assembly against Genbank |
| 28/08/2013 | Rachel Glover | Blastx of KW1 assembly against Genbank |
| 28/08/2013 | Rachel Glover | RNAseq Chalara fraxinea seqs: FERA_94 includes reads trinity assembly blastx of the assembly against genbank |
| 28/08/2013 | Rachel Glover | RNAseq Chalara fraxinea seqs: FERA_93 includes reads trinity assembly blastx of the assembly against genbank |
| 28/08/2013 | Rachel Glover | RNAseq Chalara fraxinea seqs: FERA_88 includes reads trinity assembly blastx of the assembly against genbank |
| 28/08/2013 | Rachel Glover | RNAseq Chalara fraxinea seqs: FERA_233 includes reads trinity assembly blastx of the assembly against Genbank |
| 28/08/2013 | Rachel Glover | RNAseq Chalara fraxinea seqs: FERA_232 includes reads trinity assembly blastx of the assembly against genbank |
| 28/08/2013 | Rachel Glover | RNAseq Chalara fraxinea seqs: FERA_105 includes reads trinity assembly blastx of the assembly against genbank |
| 14/08/2013 | Martin Trick | Added Tree35 RNA-seq reads assemblies annotation and BLAST results |

| | | |
|---|---|---|
| 09/08/2013 | ethering | Added GIRM2 and LSVM82 reads and analysis |
| 12/07/2013 | Jason Stajich | re-ran with multiple input seqs so the coords showup properly |
| 08/07/2013 | Jason Stajich | annotation description for antiSMASH |
| 08/07/2013 | Jason Stajich | annotation table for making EMBL file |
| 08/07/2013 | Jason Stajich | some scripts for prepping and running antismash |
| 04/07/2013 | Diane Saunders | mRNAseq data and assemblies of primordial and mature fruiting bodies |
| 17/06/2013 | Rachel Glover | assembly of the Upton broad and marshes RNA-seq data |
| 17/06/2013 | Rachel Glover | assembly of the Holt Country Park RNA-seq data |
| 31/05/2013 | Jason Stajich | added LTRharvest output |
| 14/05/2013 | Dan MacLean | Merge pull request #5 from bjclavijo/master Nornex Tree35 assembled by TGAC |
| 13/05/2013 | Bernardo J. Clavijo (TGAC) | Added Nornex tree35 assembly by TGAC |
| 10/05/2013 | Diane Saunders | Expression analysis of Ash genes |
| 01/05/2013 | Dan MacLean | add SNP/indel calls for AT1 AT2 UB1 vs TGAC 1 SAMtools called SNP/indels in VCF format and BAM files for 3 read sets vs TGAC 1 assembly |
| 26/04/2013 | Diane Saunders | Added sequences from Primordial fruit bodies |
| 18/04/2013 | Richard Smith | Fix 2nd reads filename |
| 27/03/2013 | Diane Saunders | Added trinity assembly of RNAseq data from KW1 mycelia |
| 20/03/2013 | Dan MacLean | Added Forest Research strain and mating type information Forest Research provided data on strains in their collection they have done PCR on to find out mating types. Data from Gavin Hunter and Steven Hendry. Added to repo by DM. |
| 20/02/2013 | Diane Saunders | Secretome prediction for Chalara fraxinea isolate KW1 |
| 17/02/2013 | David Swarbreck | TGAC gene predictions and source files added version Chalara_fraxinea_ass_s1v1_ann_v1.1 |
| 15/02/2013 | ethering | Updated geospatial data. Yellow pin-markers mean unlikely or unchecked data |
| 15/02/2013 | ethering | Added Geospatial data (provided by ashtag). There is also a kml file in here for loading into Google Maps |
| 13/02/2013 | ethering | Updated directory structure to provide more information about type and origin of data. Added RNAseq reads for KW1, UB1 and HP1 |
| 11/02/2013 | Diane Saunders | Added read information for RNAseq data from Upton Broad and Marshes Holt Country Park and from mycelia of KW1. |
| 28/01/2013 | ethering | Added TopHat alignment of AT1 reads against KW1 genome assembly. |
| 28/01/2013 | Diane Saunders | Added MAT locus analysis of AT1 and AT2 assembled transcripts |
| 28/01/2013 | Diane Saunders | Added Trinity assembly of ATU1 and blast against eukaryotic database |
| 26/01/2013 | Diane Saunders | Added alignment.info file for alignment of AT1 reads against KW1 assembly |
| 25/01/2013 | Diane Saunders | tblastn with MAT locus elements |
| 24/01/2013 | Dan MacLean | updated assembly.info for chalara KW1 assembly |
| 22/01/2013 | Diane Saunders | Added KW1 genome assembly version 1 |
| 18/01/2013 | Dan MacLean | Added Ash reads file File of ftp-site location for ash reads |
| 16/01/2013 | Diane Saunders | Modified file structure and added metadata for uninfected Ash sample |
| 24/12/2012 | BPatrickChapman | added signalP analysis of AT1 likely coding sequences |
| 21/12/2012 | MattBashton | added AT2_SF_run_with_pngs.tbz2 |
| 21/12/2012 | MattBashton | Added Pfam A Vs AT2 Trinity-r2012-10-05 assembly likely coding sequences |
| 21/12/2012 | MattBashton | Added alternative assembly for AT1 produced with Trinity-r2012-10-05 |
| 19/12/2012 | Dan MacLean | fixed typo in git url in readme |
| 19/12/2012 | Diane Saunders | AT2 trinity assembly version 2 blastx against eukaryotic database |
| 19/12/2012 | Diane Saunders | Added alternative AT2 assembly using Trinity |
| 19/12/2012 | MattBashton | added SUPERFAMILY domain assignments for AT2 likely coding sequences |
| 19/12/2012 | MattBashton | Merge branch 'master' of https://github.com/ash-dieback-crowdsource/data |

| | | |
|---|---|---|
| 19/12/2012 | MattBashton | added Helotiales NPP alignment of 84 sequences |
| 19/12/2012 | MattBashton | added AT1 and AT2 NPP domains aligned to Pfam seed |
| 19/12/2012 | MattBashton | added AT2 NPP domains |
| 19/12/2012 | MattBashton | added AT2 likely_coding_sequences from trinity |
| 18/12/2012 | MattBashton | added AT2 assembly from trinity |
| 17/12/2012 | MattBashton | added model.tab and dir.cla.scop.txt to allow cross-ref of IDs in Ash.ass |
| 17/12/2012 | MattBashton | added alignment of AT1 NPP sequences to those in NPP1 Pfam seed |
| 17/12/2012 | MattBashton | Merge branch 'master' of https://github.com/ash-dieback-crowdsource/data |
| 17/12/2012 | MattBashton | added Pfam A Vs AT1 CDS |
| 17/12/2012 | Crossmanlc | predicted_pfam_map_to_GO |
| 17/12/2012 | MattBashton | added AT1_SF_run_with_pngs.tbz2Âand updated .info |
| 17/12/2012 | MattBashton | SUPERFAMILY domain assignments for AT1 likely coding sequences |
| 17/12/2012 | MattBashton | Elegant and fast way to find NPP domains |
| 17/12/2012 | MattBashton | Likely Coding Sequences |
| 17/12/2012 | MattBashton | FastQC on reads |
| 16/12/2012 | Sophien Kamoun | BLASTN of 116 Chalara fraxinea sequences in GenBank vs AT1 assembly |
| 16/12/2012 | Sophien Kamoun | polyketide synthase comp1171 amino acid sequence |
| 16/12/2012 | Sophien Kamoun | LysM effector comp8971 amino acid sequence |
| 14/12/2012 | Diane Saunders | added AT1_NLP1.fan |
| 14/12/2012 | Diane Saunders | Details of NRP-like transcripts added |
| 13/12/2012 | Kentaro Yoshida | Add blast folder to ashwellthorpe AT1 Add blast folder to ashwellthorpe AT1 |
| 13/12/2012 | Kentaro Yoshida | Add BLASTP output of custom eukaryote protein database Add BLASTP output of custom eukaryote protein database Diane analyzed |
| 13/12/2012 | Kentaro Yoshida | Add TBLATN output of pks Add TBLATN output of pks Sophien analyzed |
| 13/12/2012 | Kentaro Yoshida | Add TBLASTN of NLPs Add TBLASTN of NLPs Sophien analyzed |
| 13/12/2012 | Kentaro Yoshida | Add TBLASTN output of LysM Add TBLASTN output of LysM Sophien analyzed |
| 12/12/2012 | Kentaro Yoshida | Upload AT1 assembly Upload AT1 assembly |
| 11/12/2012 | Dan MacLean | TSL/JIC first data added Some Calmodulin and ITS sequence from the Ashwellthorpe Wood and Kenninghall Wood branch material added. Also added RNAseq reads from same. |
| 16/11/2012 | Dan MacLean | initial push |

Table S1: All commits to the OADB GitHub repository. Full text version available at `http://dx.doi.org/10.6084/m9.figshare.1005035`.

| GO Term | Description | Secretome Genes | Non-Secretome Genes | Proportion Secretome | Combined Probability | Population | Bonferroni Corrected p-value |
|---|---|---|---|---|---|---|---|
| GO:0016798 | hydrolase activity, acting on glycosyl bonds | 1 | 0 | 1 | 0.06 | 1 | 0 |
| GO:0046558 | arabinan endo-1,5-alpha-L-arabinosidase activity | 1 | 0 | 1 | 0.06 | 1 | 0 |
| GO:0016901 | oxidoreductase activity, acting on the CH-OH group of donors, quinone or similar compound as acceptor | 1 | 0 | 1 | 0.06 | 1 | 0 |
| GO:0005199 | structural constituent of cell wall | 2 | 0 | 1 | 0.00 | 2 | 0 |
| GO:0006078 | (1-6)-beta-D-glucan biosynthetic process | 1 | 0 | 1 | 0.06 | 1 | 0 |
| GO:0004867 | serine-type endopeptidase inhibitor activity | 1 | 0 | 1 | 0.06 | 1 | 0 |
| GO:0005975 | carbohydrate metabolic process | 121 | 116 | 0.51 | 0 | 237 | 0 |
| GO:0004557 | alpha-galactosidase activity | 2 | 0 | 1 | 0.00 | 2 | 0 |
| GO:0005576 | extracellular region | 22 | 3 | 0.88 | 0 | 25 | 0 |
| GO:0071555 | cell wall organization | 1 | 0 | 1 | 0.06 | 1 | 0 |
| GO:0052861 | glucan endo-1,3-beta-glucanase activity, C-3 substituted reducing group | 1 | 0 | 1 | 0.06 | 1 | 0 |
| GO:0003980 | UDP-glucose:glycoprotein glucosyltransferase activity | 1 | 0 | 1 | 0.06 | 1 | 0 |
| GO:0016158 | 3-phytase activity | 1 | 0 | 1 | 0.06 | 1 | 0 |
| GO:0006308 | DNA catabolic process | 2 | 0 | 1 | 0.00 | 2 | 0 |
| GO:0006004 | fucose metabolic process | 1 | 0 | 1 | 0.06 | 1 | 0 |
| GO:0030570 | pectate lyase activity | 2 | 0 | 1 | 0.00 | 2 | 0 |
| GO:0016671 | oxidoreductase activity, acting on a sulfur group of donors, disulfide as acceptor | 1 | 0 | 1 | 0.06 | 1 | 0 |
| GO:0000298 | endopolyphosphatase activity | 1 | 0 | 1 | 0.06 | 1 | 0 |
| GO:0030245 | cellulose catabolic process | 2 | 0 | 1 | 0.00 | 2 | 0 |
| GO:0003756 | protein disulfide isomerase activity | 1 | 0 | 1 | 0.06 | 1 | 0 |
| GO:0005773 | vacuole | 2 | 0 | 1 | 0.00 | 2 | 0 |
| GO:0052862 | glucan endo-1,4-beta-glucanase activity, C-3 substituted reducing group | 1 | 0 | 1 | 0.06 | 1 | 0 |
| GO:0009277 | fungal-type cell wall | 1 | 0 | 1 | 0.06 | 1 | 0 |
| GO:0019028 | viral capsid | 2 | 0 | 1 | 0.00 | 2 | 0 |
| GO:0004553 | hydrolase activity, hydrolyzing O-glycosyl compounds" | 87 | 54 | 0.62 | 0 | 141 | 0 |
| GO:0008810 | cellulase activity | 2 | 0 | 1 | 0.00 | 2 | 0 |
| GO:0043086 | negative regulation of catalytic activity | 4 | 0 | 1 | 0.00 | 4 | 0 |
| GO:0004556 | alpha-amylase activity | 1 | 0 | 1 | 0.06 | 1 | 0 |
| GO:0004563 | beta-N-acetylhexosaminidase activity | 3 | 0 | 1 | 0.00 | 3 | 0 |
| GO:0031221 | arabinan metabolic process | 1 | 0 | 1 | 0.06 | 1 | 0 |
| GO:0004560 | alpha-L-fucosidase activity | 3 | 0 | 1 | 0.00 | 3 | 0 |

| | | | | | | | |
|---|---|---|---|---|---|---|---|
| GO:0045735 | nutrient reservoir activity | 6 | 0 | 1 | 0 | 6 | 0 |
| GO:0006508 | proteolysis | 54 | 96 | 0.36 | 0 | 150 | 0 |
| GO:0008474 | palmitoyl-(protein) hydrolase activity | 1 | 0 | 1 | 0.06 | 1 | 0 |
| GO:0015926 | glucosidase activity | 1 | 0 | 1 | 0.06 | 1 | 0 |
| GO:0006491 | N-glycan processing | 1 | 0 | 1 | 0.06 | 1 | 0 |
| GO:0004521 | endoribonuclease activity | 1 | 0 | 1 | 0.06 | 1 | 0 |
| GO:0003824 | catalytic activity | 123 | 556 | 0.18 | 0 | 679 | 1.00E-10 |
| GO:0016614 | oxidoreductase activity, acting on CH-OH group of donors | 32 | 63 | 0.34 | 0 | 95 | 1.33E-08 |
| GO:0008236 | serine-type peptidase activity | 15 | 12 | 0.56 | 0 | 27 | 2.43E-07 |
| GO:0004252 | serine-type endopeptidase activity | 17 | 20 | 0.46 | 0 | 37 | 1.34E-06 |
| GO:0008061 | chitin binding | 9 | 3 | 0.75 | 0 | 12 | 4.56E-06 |
| GO:0004650 | polygalacturonase activity | 9 | 3 | 0.75 | 0 | 12 | 4.56E-06 |
| GO:0004601 | peroxidase activity | 11 | 8 | 0.58 | 0 | 19 | 1.90E-05 |
| GO:0005618 | cell wall | 8 | 3 | 0.73 | 0 | 11 | 4.12E-05 |
| GO:0050660 | flavin adenine dinucleotide binding | 34 | 106 | 0.24 | 0 | 140 | 6.67E-05 |
| GO:0050525 | cutinase activity | 6 | 1 | 0.86 | 0 | 7 | 0.00 |
| GO:0030246 | carbohydrate binding | 15 | 23 | 0.39 | 0 | 38 | 0.00 |
| GO:0030248 | cellulose binding | 6 | 2 | 0.75 | 0 | 8 | 0.00 |
| GO:0016052 | carbohydrate catabolic process | 9 | 8 | 0.529411765 | 0 | 17 | 0.00 |
| GO:0042545 | cell wall modification | 5 | 1 | 0.83 | 0.00 | 6 | 0.00 |
| GO:0030599 | pectinesterase activity | 5 | 1 | 0.83 | 0.00 | 6 | 0.00 |
| GO:0008762 | UDP-N-acetylmuramate dehydrogenase activity | 16 | 35 | 0.31 | 0 | 51 | 0.00 |
| GO:0042802 | identical protein binding | 4 | 1 | 0.8 | 0.00 | 5 | 0.01 |
| GO:0016788 | hydrolase activity, acting on ester bonds | 12 | 25 | 0.32 | 0 | 37 | 0.03 |
| GO:0008233 | peptidase activity | 10 | 18 | 0.36 | 0 | 28 | 0.04 |
| GO:0005507 | copper ion binding | 10 | 18 | 0.36 | 0 | 28 | 0.04 |
| GO:0004190 | aspartic-type endopeptidase activity | 9 | 15 | 0.375 | 0 | 24 | 0.05 |

Table S2: Enriched GO Terms in the secretome relative to the whole genome at Bonferroni corrected p <0.05. Full text version available at `http://dx.doi.org/10.6084/m9.figshare.1005019`
